# Metabolically robust autoluminescent reporters

**DOI:** 10.1101/2025.08.04.668150

**Authors:** Viktor V Morozov, Anastasia V Balakireva, Feng Gao, Aubin Fleiss, Nataliya M Mishina, Maxim M Perfilov, Tatiana Yu Mitiouchkina, Dmitry A Gorbachev, Keith V Wood, Ilia V Yampolsky, Karen S Sarkisyan, Alexander S Mishin

## Abstract

Self-sustained luminescence can be engineered in non-luminous organisms by integrating luciferin biosynthesis pathways with the host’s metabolism. When conditionally expressed, such pathways enable non-invasive monitoring of virtually any transcriptional event, providing a wide dynamic range and high resolution. However, because light emission depends on substrate availability within the cell, autoluminescence reporters are inherently non-quantitative: signal intensity reflects both gene expression and cellular metabolism. Here, we present an approach that disentangles the contributions of gene expression and metabolism, rendering autoluminescence signals robust against metabolic perturbations. By co-expressing an engineered reference luciferase variant that uses the same substrate but emits light at a different wavelength, and monitoring the resulting colour ratio, we dramatically reduce signal variation, enabling quantitative physiological imaging amid changing metabolic activity. Our strategy is applicable across known autoluminescence pathways from bacteria and fungi.

---

Luminescence is commonly used to measure gene expression. Typically, a luciferase-expressing cell is assayed in the presence of excess of the substrate, luciferin. When no cofactor is required, light intensity reflects the amount of active luciferase, which is used as a proxy for gene expression. In the absence of recombinant luciferase, most organisms emit virtually no light — the reason why luminescence imaging is often favoured, as very low background allows for high signal-to-noise ratio and wide dynamic range. But at the whole-organism scale, approaches that rely on exogenously supplied substrates suffer from invasiveness, high cost, poor uptake, uneven luciferin distribution, and sometimes, toxicity ^1–3^.

An alternative way to achieve luminescence is through intracellular production of substrate. Currently, two pathways are understood enough to allow heterologous luciferin biosynthesis **(Figure 1a)**; these pathways originate from bacteria and fungi, and both have been adapted as a biochemical basis for engineering autoluminescence ^4–8^. Bacterial pathway, a branch of fatty acid metabolism, found most use in bacterial hosts and animal cell culture ^9,10^. The fungal pathway, a branch of phenylpropanoid metabolism, was shown to be easily integratable into the metabolism of plants and fungi ^11–16^.

**Figure 1.**
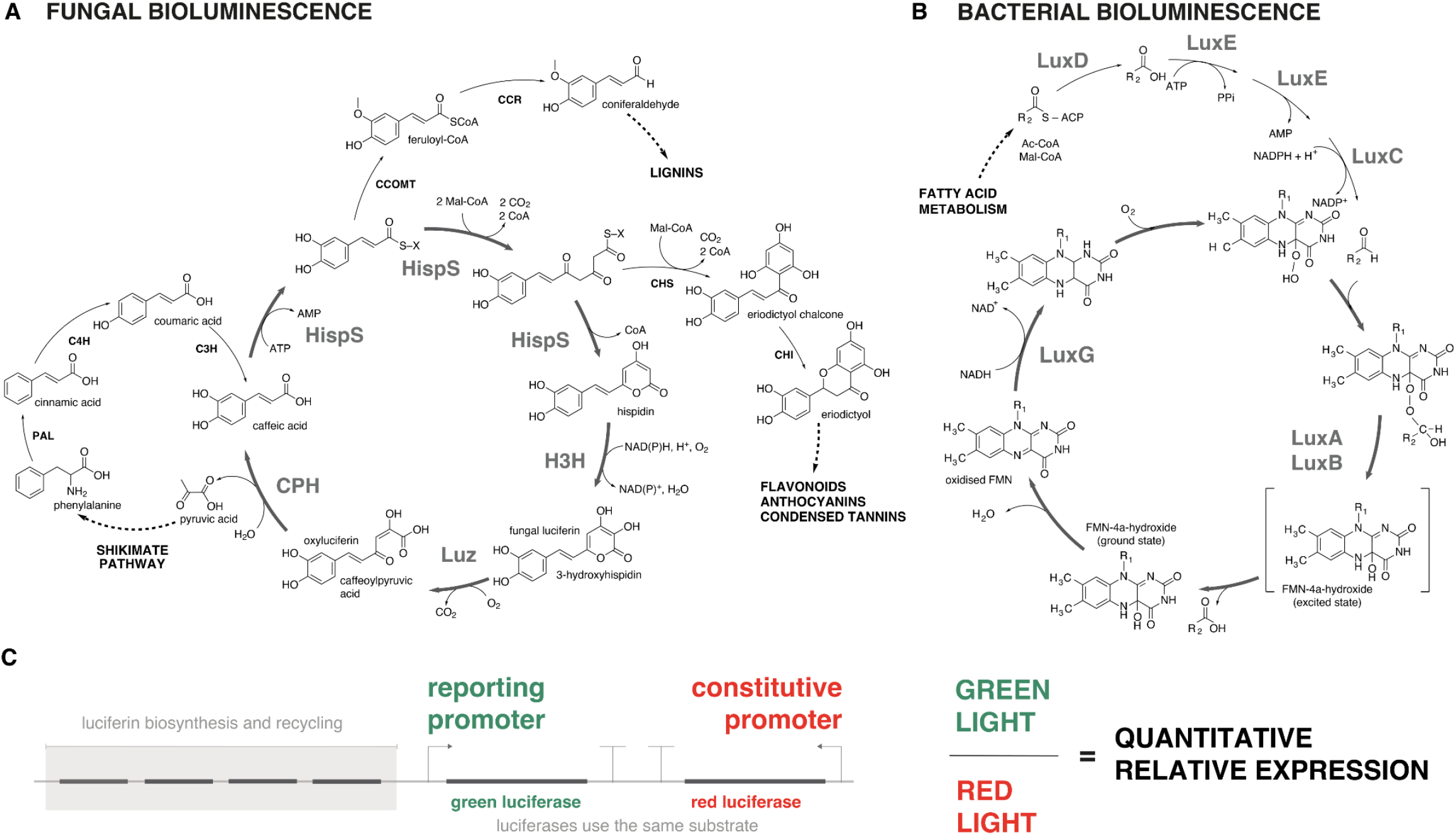
Bioluminescent pathways in fungi and bacteria, and dual-colour reporter design. **A**. Schematic of the fungal bioluminescence pathway in the context of plant metabolism. Shikimate and phenylpropanoid pathways lead to biosynthesis of caffeic acid, which is subsequently used to generate hispidin and fungal luciferin. In bioluminescent plants, key bioluminescence-specific enzymes such as Hisps (hispidin synthase), H3H (hispidin-3-hydroxylase), Luz (fungal luciferase), and CPH (caffeoylpyruvate hydrolase) are expressed. Competing branches lead to lignin and flavonoid biosynthesis. **B.** Overview of the bacterial bioluminescence pathway. The fatty acid metabolism-derived aldehyde is processed through the LuxCDABE pathway. LuxCDE enzymes generate the long-chain fatty aldehyde substrate; LuxAB encodes luciferase; LuxG supplies reduced flavin mononucleotide (FMNH_2_) required for light production. C. Design of a dual-luciferase expression reporter. Genes encoding green and red luciferases (which use the same luciferin substrate) are placed under separate regulatory elements: a constitutive promoter (red) and a variable, reporting promoter (green). Light registered at each wavelength enables ratiometric quantification of promoter activity.

However, all self-sustained luminescence technologies are inherently non-quantitative due to reliance on metabolic activity. Luciferin is produced biosynthetically in every cell, and light emission reflects not only the expression of bioluminescence genes but also the metabolic activity, including production and uptake of biochemical precursors, reducing equivalents and energy supply ^17^. In addition, environmental factors, such as temperature or oxygen availability, influence light emission, interfering with quantitative imaging ^18^.

Here, we aimed to engineer autoluminescence systems to report gene expression independently of metabolism. If two luciferases that use the same substrate but emit light of different colours are co-expressed, the colour ratio should reflect the relative abundance of these luciferases and be largely independent of substrate concentration (Figure 1c).

We first focused on creating a version of the fungal bioluminescence pathway that emits red-shifted light by engineering resonance energy transfer between luciferase and a fluorescent protein ^19^. To identify an optimal fusion topology, we designed an experiment to probe different potential insertion points. Starting with a gene encoding nnLuz_v4, an improved version of *Neonothopanus nambi* luciferase ^14^, we created a library of variants where a DNA insert was sliding along the sequence, being introduced in frame at every ninth nucleotide of the gene (**Supplementary Figure 1a**). This insert sequence was designed to (1) encode 17 amino acids, and expected to fold into a flexible loop, (2) contain two recognition sites for MlyI endonuclease to allow generation of a 2-aa deletion library, (3) contain two recognition sites for BtgZI to replace the insert with another protein domain, leaving only 1-aa scars. Screening of the insertion and deletion libraries in plant cells revealed positions tolerant to structural perturbations. As expected, these mostly included residues in protein termini. However, we also found positions within the protein sequence where insertions and deletions resulted in functional proteins (**Supplementary Figure 1b, Figure 2a**).

**Figure 2.**
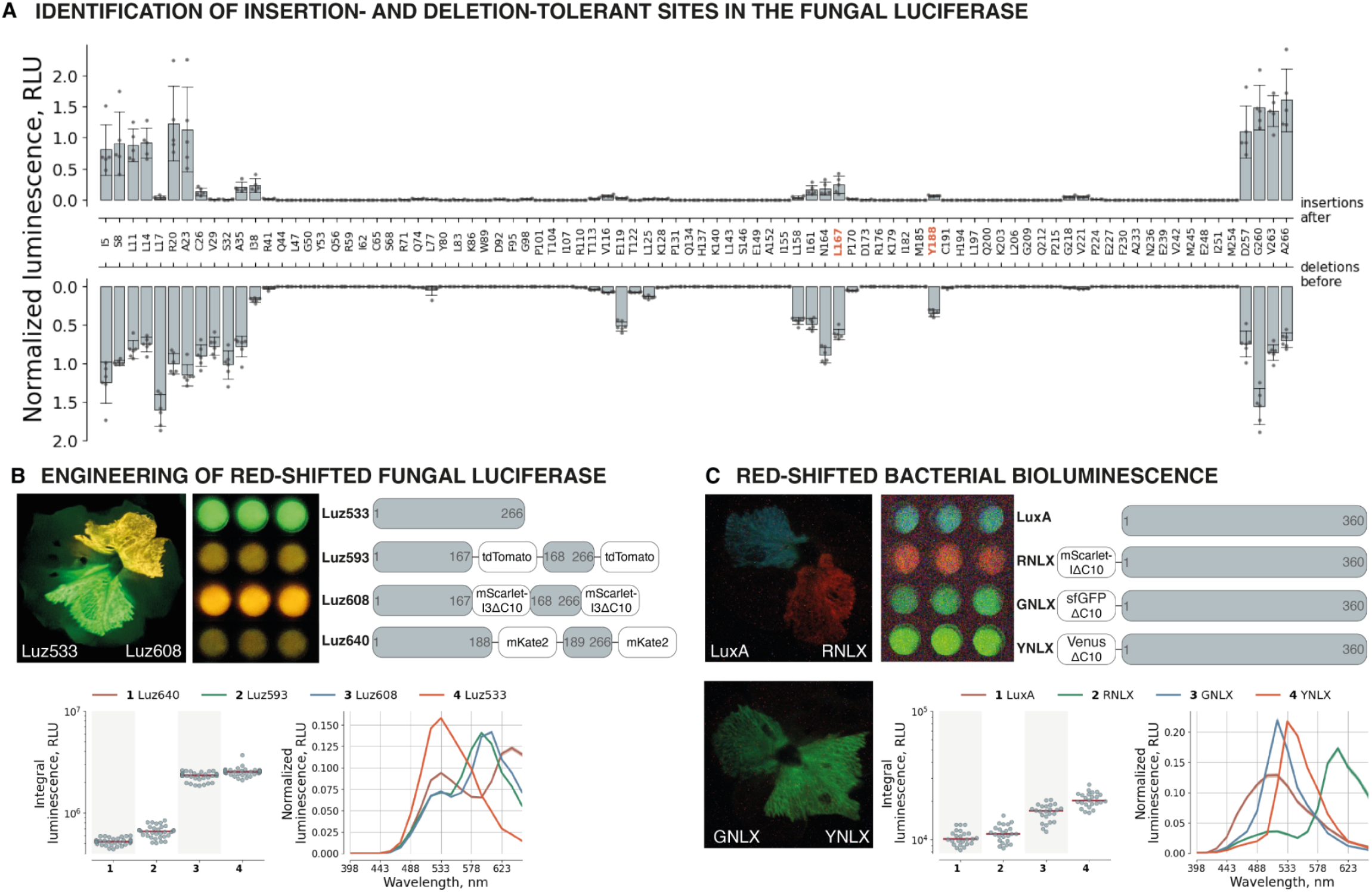
Engineering of red-shifted fungal luciferase. **A.** Effect of a 17-amino-acid sequence insertion or 2-amino-acid deletions in fungal luciferase expressed in tobacco BY-2 plant cell packs. Insertion and deletion sites are shown on the X-axis. Data points represent average luminescence, normalized to the luminescence of the “wild type” nnLuz_v4, N = 6 **B.** Images of *Petunia hybrida* flowers (shot on iPhone 15 Pro) and BY-2 cell packs (shot on Sony alpha), protein structures, brightness, and emission spectra of selected fungal luciferase variants: nnLuz_v5 (Luz533), and its fusions with tdTomato (Luz593), mScarlet-I3ΔC10 (Luz608), and mKate2 (Luz640). Median values are indicated by red lines in the swarm plots. All the values are depicted in the swarm as grey dots. The spectra are presented as mean values normalised to the sum of luminescence intensities at every wavelength and standard deviation, N = 30. **C.** Images of *Petunia hybrida* flowers (shot on Sony alpha), BY-2 cell packs, protein structures, brightness, and emission spectra of color-shifted bacterial bioluminescence pathway variants from Kusuma et al. ^5^ Median values are indicated by red lines in the swarm plots. All the values are depicted in the swarm as grey dots. The spectra are presented as mean values normalised to the sum of luminescence intensities at every wavelength and standard deviation, N = 24.

We chose four of the identified internal regions — E119, I161-L167, Y188, and V221 — to insert different red fluorescent proteins and assess BRET efficiency and brightness in plant cells (**Supplementary Figure 2**). Two variants, nnLuz167-tdTomato and nnLuz188-mKate2, showed significantly red-shifted spectra, but were dim. By optimising linker composition and length, replacing luciferase with an improved version, nnLuz_v5, obtained in a parallel study, and attaching an additional fluorescent protein to the termini (**Supplementary Figures 3-8, Supplementary Tables 1-5**), we identified three variants with optimised brightness and efficient resonance transfer (**Figure 2b**), which we called Luz593, Luz608 and Luz640. When expressed in *Nicotiana tabacum* BY-2 cells or in the petals of *Petunia hybrida*, all variants showed colour changes easily detectable on regular consumer cameras without filters (**Figure 2b**). Although the spectral separation was the largest for Luz640, we chose Luz593 and Luz608 for future experiments because of their higher brightness.

**Figure 3.**
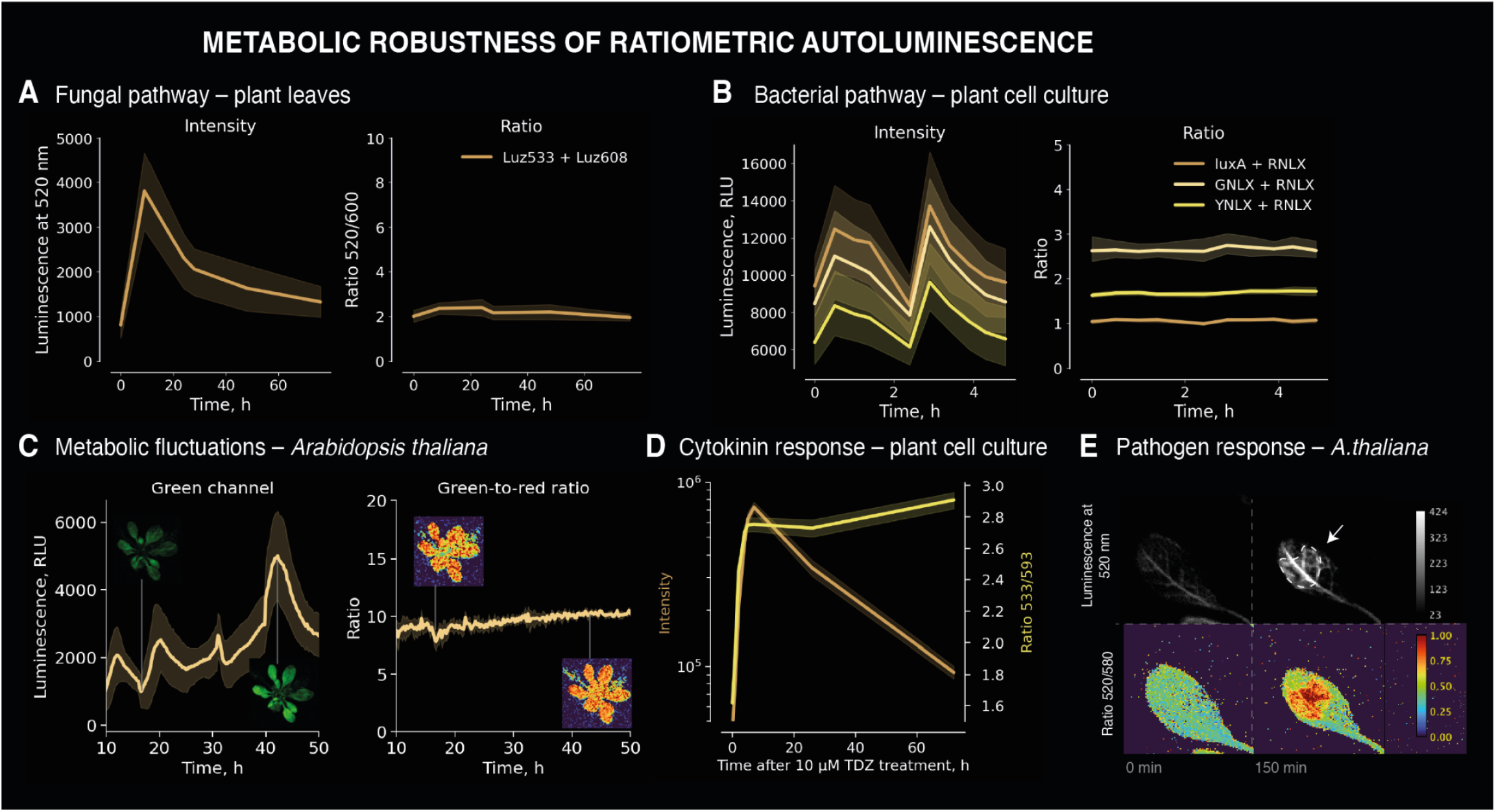
Metabolic robustness of ratiometric autoluminescence. Tolerance of the ratiometric bioluminescence readout to metabolic perturbations: **A.** *Nicotiana benthamiana* leaves, fungal bioluminescence pathway, treated with 1 mM solution of thidiazuron, N = 4; **B.** *Nicotiana tabacum* BY-2 cell packs, bacterial bioluminescence pathway, submerged into an infiltration buffer to alter oxygen availability, N = 8; **C.** Transgenic *Arabidopsis thaliana*, ratiometric salicylic acid reporter, fungal bioluminescence, no treatments, N = 5. D. Response of the fungal bioluminescence-based ratiometric cytokinin reporter to treatment with 10 µM thidiazuron, N = 24. E. The response of the fungal bioluminescence-based ratiometric jasmonic acid reporter to the *Pseudomonas savastanoi* infiltration in the green channel (520 nm) and in ratio 520/580. All plots show mean values and standard deviation.

We then used these variants to test whether the ratiometric autoluminescence imaging can reduce dependence of reporters readout on metabolic activity. First, as a proof-of-concept, we co-expressed two luciferases of different colours, controlled by strong constitutive promoters. Co-expression of red Luz608 and green Luz533 in the plant cell culture (**Supplementary Figures 10**) or *Nicotiana benthamiana* leaves (**Figure 3a, Supplementary Figures 11-12**), led to a major stabilisation of the signal, compared to intensity-based measurements.

Similarly, we performed an experiment with the metabolically distinct bacterial bioluminescence pathway. We cloned the colour-shifted variants of the blue light-emitting bacterial luciferase LuxA ^5^: green (GNLX = sfGFPΔC10_EL_iluxA), yellow (YNLX = VenusΔC10_EL_iluxA), and red (RNLX = mScarlet_IΔC10_EL_iluxA), and expressed them in the plant cell culture (**Figure 2c**). We observed a similar effect: the ratiometric readout was up to 16-fold more stable than intensity-based measurements (**Figure 3b, Supplementary Figure 9**).

Next, we tested whether the ratiometric approach allows disentangling of metabolic perturbations from gene expression changes when the reporter is driven by physiologically relevant promoters. As the bacterial pathway produced ∼2 orders of magnitude less light than the fungal pathway in our hands, we chose the fungal pathway for further work. In our next experiment in plant cell culture, the cytokinin-sensitive promoter pTCSv2 [ref ^20^] drove the expression of green Luz533, while the reference red Luz593 was controlled by the constitutive promoter 35S. In the green channel, after reaching the maximum, luminescence dropped almost 10-fold over time, while the ratiometric signal remained largely stable (**Supplementary Figure 10**). In contrast, when treated with cytokinin, the ratio showed expected dose-dependent response (**Figure 3d, Supplementary Figure 10**), providing quantitative readout amid metabolic fluctuations. Similar results were obtained upon agroinfiltration of *Nicotiana benthamiana* leaves (**Supplementary Figures 11-12**), where we used Luz608 as the red luciferase. In this case, we additionally confirmed with qPCR that observed ratio changes correspond to changes in gene expression (**Supplementary Figures 12d**).

Finally, we created transgenic *Arabidopsis thaliana* plants. We used hormone-sensitive promoters for the green luciferase nnLuz_v5 (salicylic-acid-sensitive pWRKY70 ^21^ or jasmonic-acid–sensitive pORCA3 promoter ^22^), while the red luciferase Luz593 was expressed from the constitutive promoter pMinSyn108 ^23^ (**Supplementary Table 6**). Life-long luminescence imaging of the salicylic-acid-reporting line (**Supplementary Figure 13, Supplementary Video 1**) showed that despite the fluctuations in luminescence intensity the ratio of the signals was stable in the absence of salicylic-acid-inducing stimuli (**Figure 3c**). In agreement with previous reports, in rosette leaves we observed flickering patterns of salicylic acid activity (**Supplementary Figure 13**) ^24^. On the jasmonic-acid-reporting line, we modeled hormonal response to an infection with *Pseudomonas savastanoi* (**Supplementary Figure 14, Supplementary videos 3-4**). Similarly, we observed that the ratiometric measurement remained stable despite multifold variations in overall luminescence intensity. This intrinsic robustness revealed the area of local response to the pathogen around the infection site, amid a highly varied background of bright leaf vasculature (**Figure 3e**).

Overall, our work shows that by co-expressing two spectrally diverse luciferases that utilise the same substrate, it is possible to convert raw autoluminescence signal into a ratiometric readout that is robust to metabolic fluctuations. Across fungal and bacterial pathways, in plant cells and whole plants, this approach dramatically reduces signal variation and allows monitoring of hormone-responsive promoter activity amid varying tissue metabolism.

Dual-colour autoluminescent constructs provide a straightforward route to quantitative imaging of molecular physiology in plants and other organisms.

## Supporting information

Supplementary Video 1.mov

## Acknowledgements

This study was partially funded by Light Bio (light.bio) and Planta (planta.bio). The Synthetic biology Group is funded by the MRC London Institute of Medical Sciences (UKRI MC-A658-5QEA0). This work was supported by UKRI Biotechnology and Biological Sciences Research Council through the International Science Partnerships Fund (ISPF) [grant number UKRI249]. Plasmid assembly was funded by RSF, project number 24-74-10105, https://rscf.ru/en/project/24-74-10105/. Luminescent assays in BY-2 cells were funded by RSF, project number 24-74-10087, https://rscf.ru/en/project/24-74-10087/. Generation of transgenic plants were funded by RSF, project number 25-76-30006, https://rscf.ru/en/project/25-76-30006/. Luminescence imaging of the transgenic plants were funded by RSF, project number 25-14-00349, https://rscf.ru/en/project/25-14-00349/.

## Data and material availability

The plasmids used in this study will be made available for non-commercial use through Addgene. Plant lines will be made available under an MTA upon request. Data is available at https://doi.org/[replace_upon_acceptance].

## Author Contributions

VVM, AVB, FG, AF, TYM, and DAG performed experimentation. VVM, AVB, AF, FG, KSS and ASM planned experiments. AVB, FG, AF, NMM, MMP, KSS and ASM performed data analysis. AVB, KSS and VVM wrote the paper. KVW, IVY, KSS and ASM, proposed and directed the study. All authors reviewed the paper draft. The authors wish it to be known that three first authors should be considered as joint first authors.

## Competing Interests

This study was partially funded by Light Bio (light.bio) and Planta (planta.bio).

