## Supplementary materials for "Metabolically robust autoluminescent reporters"

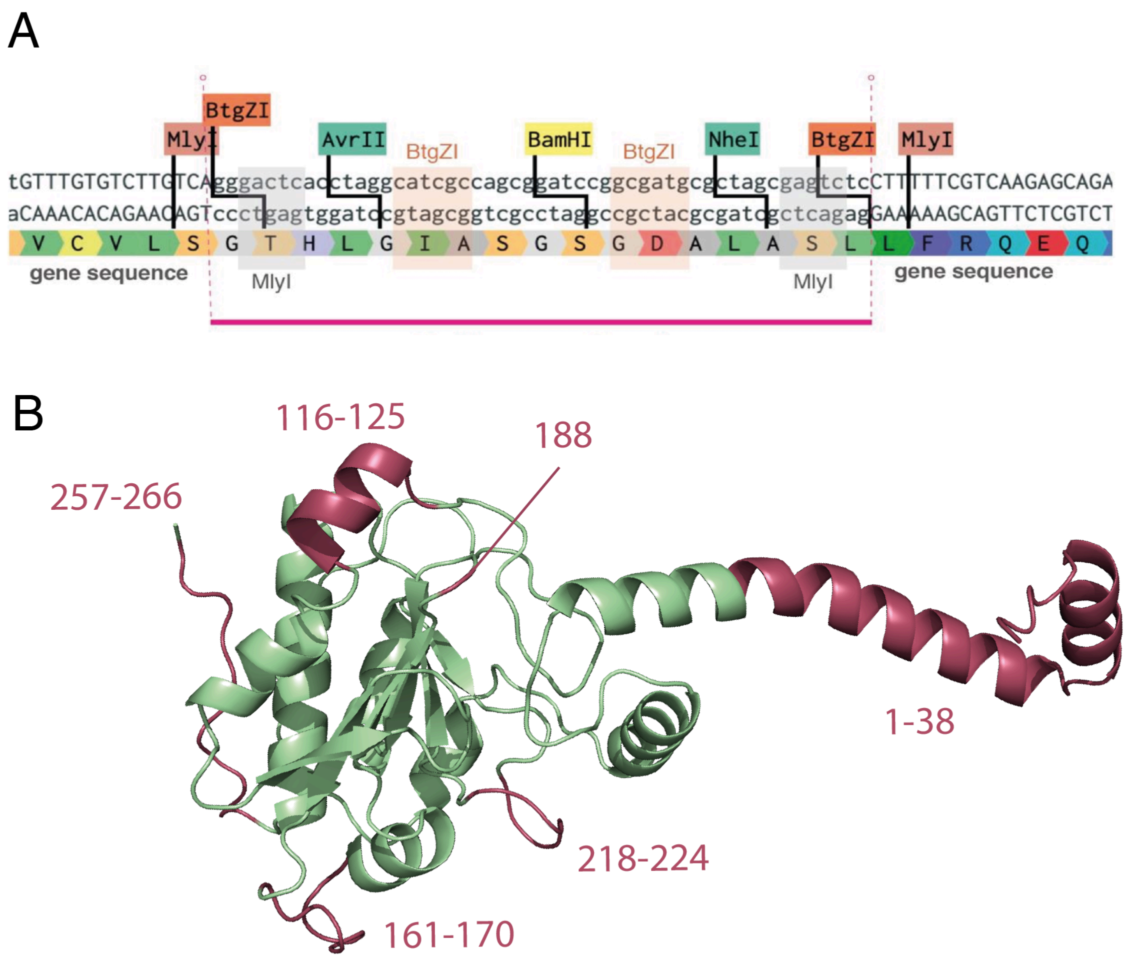


Supplementary Figure 1. Identification of sites within luciferase that are insensitive to insertions or deletions. A. Structure of the multipurpose insert in the synthetic fungal luciferase library designed for the generation of insertion and deletion libraries. B. Locations of identified positions suitable for the insertion of functional domains into fungal luciferase, indicated in pink.

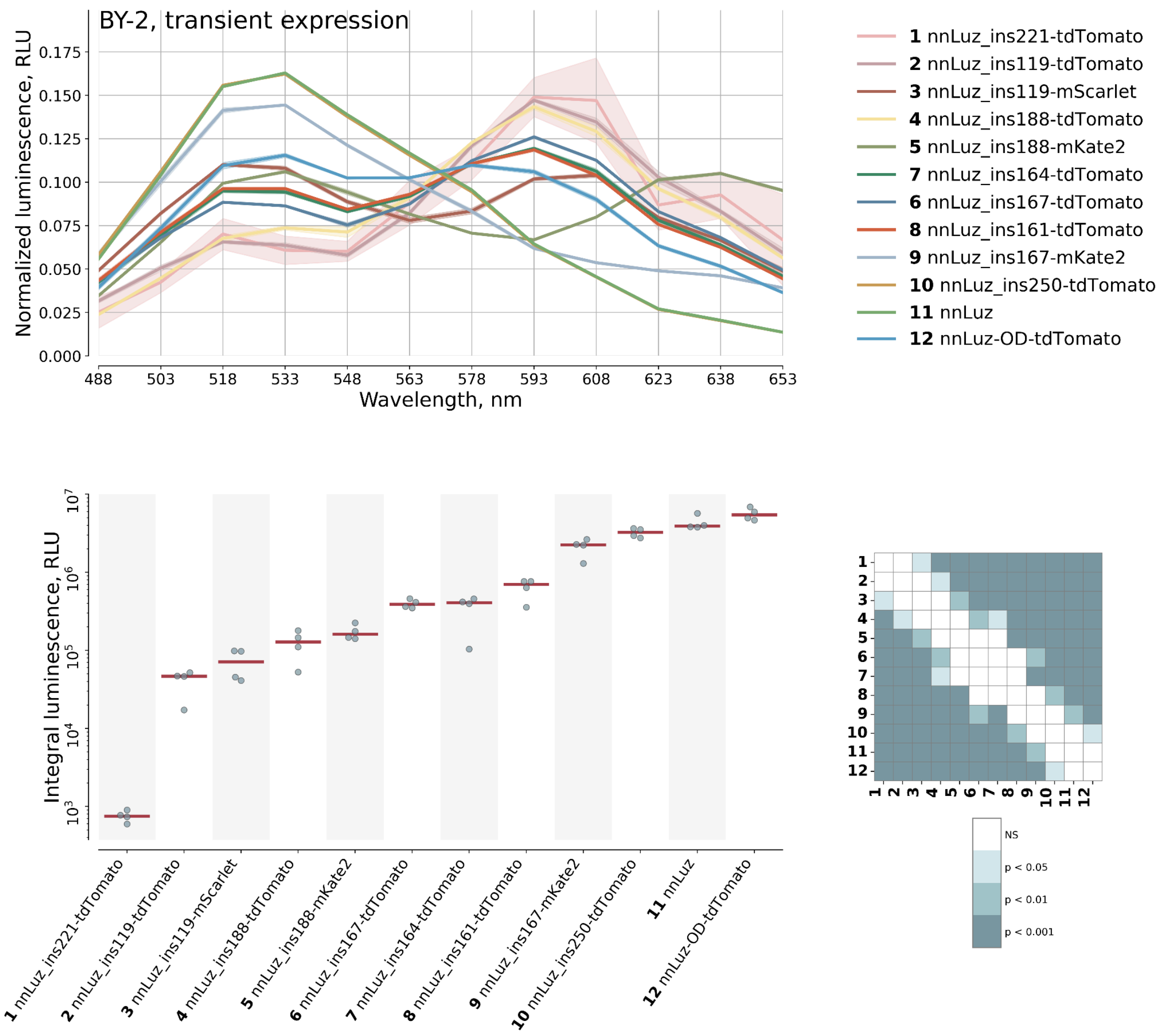


Supplementary Figure 2. **Initial screening of fluorescent proteins at the identified positions.** Spectral characteristics of the fusion proteins are shown at the top; brightness is shown at the bottom. The nnLuz label refers to the nnLuz_v4 gene variant.

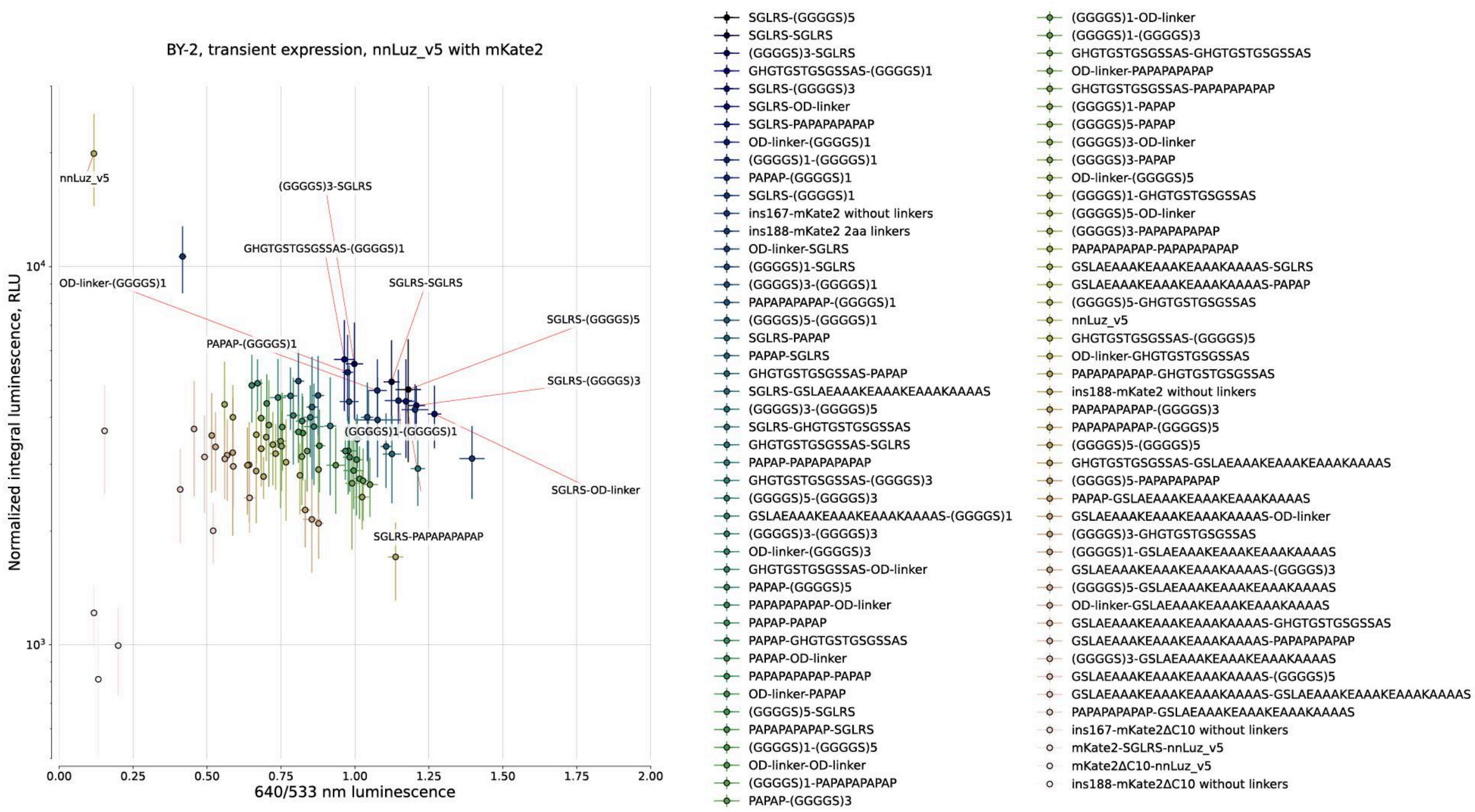


Supplementary Figure 3. Optimization of bioluminescence resonance energy transfer in the nnLuz_v5_ins188 and mKate2 pair by varying the linkers. In the legend, linker pairs are indicated as ‘N-terminal to mKate2 linker – C-terminal to mKate2 linker’. The top 10 fusions, ranked by brightness and the 640-to-533 ratio, are labeled on the plot. The raw data is indicated in Supplementary table 2.

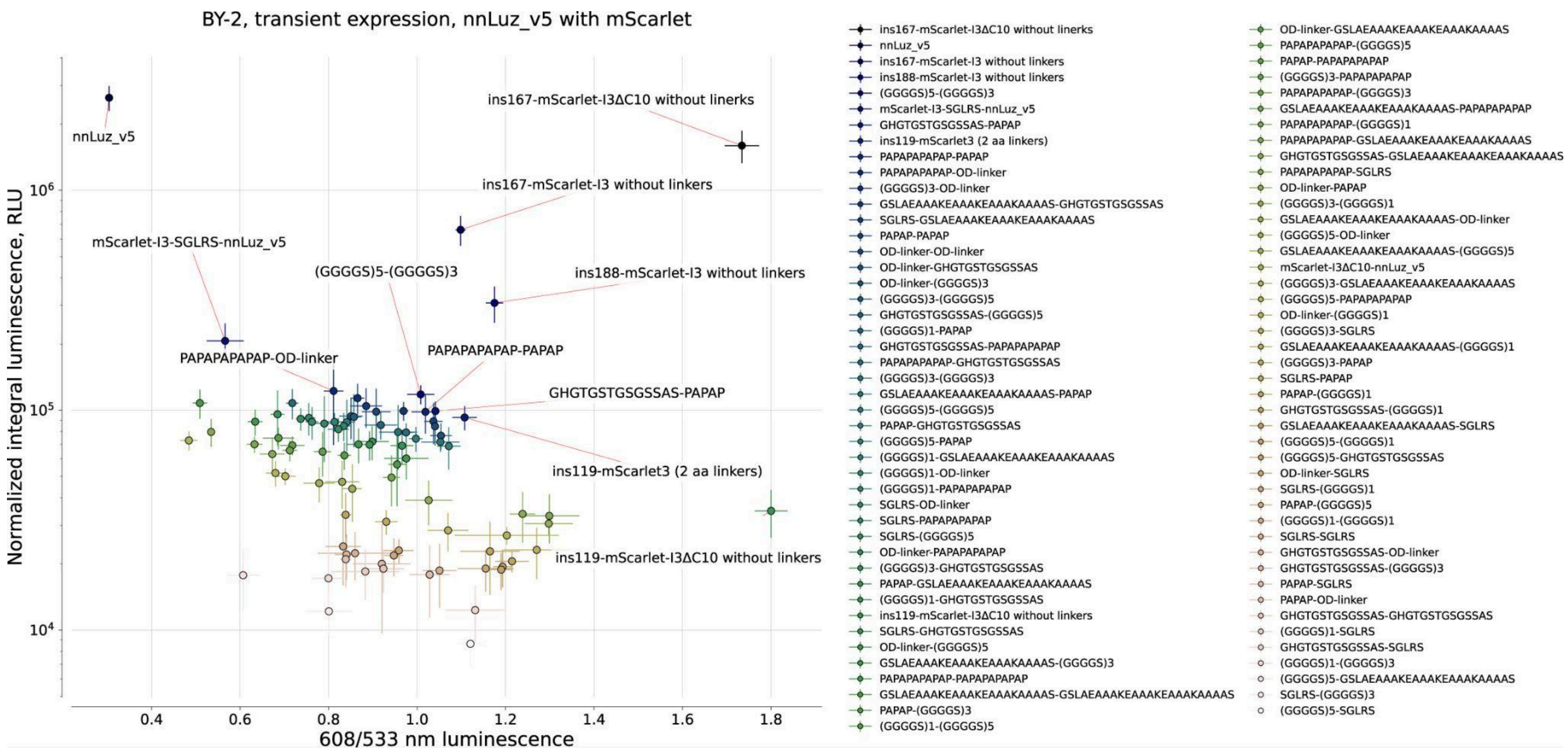


**Supplementary Figure 4. Optimization of bioluminescence resonance energy transfer in the nnLuz_v5_ins119 and mScarlet pair by varying the linkers.** In the legend, linker pairs are indicated as 'N-terminal to mScarlet linker – C-terminal to mScarlet linker'. Additionally, a comparison was conducted with a range of fusions between nnLuz_v5_ins167 or nnLuz_v5_ins188 and mScarlet-I3, designated as insX–fluorescent protein. The top 10 fusions, based on brightness and the 608-to-533 ratio, are labeled. The raw data is indicated in Supplementary table 3.

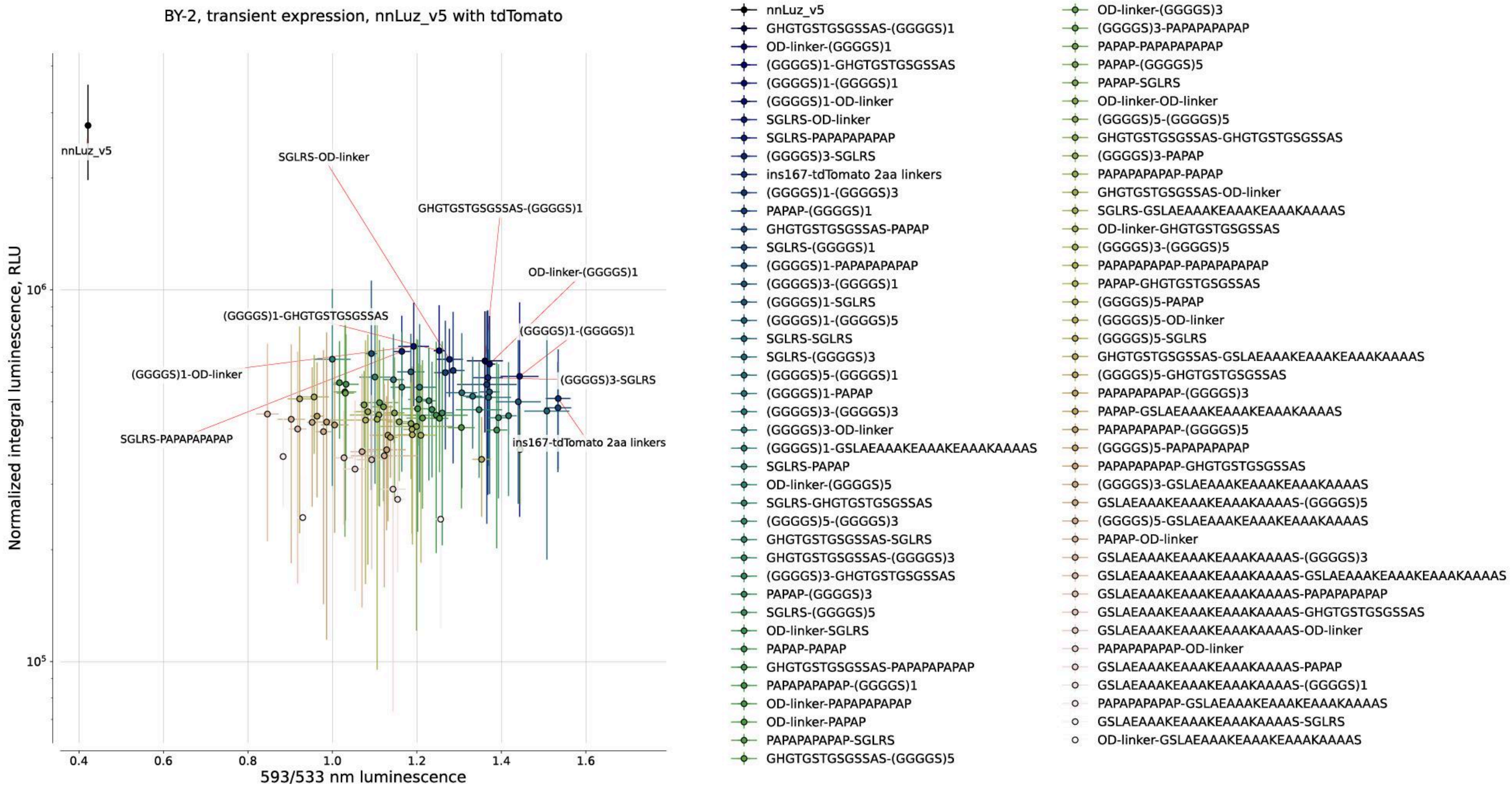


**Supplementary Figure 5. Optimization of bioluminescence resonance energy transfer in the nnLuz_v5_ins167 and tdTomato pair by varying the linkers.** In the legend, linker pairs are indicated as 'N-terminal to mScarlet linker – C-terminal to mScarlet linker'. The initial fusion of nnLuz_v5_ins167 with tdTomato, flanked by 2-amino-acid linkers, is indicated as 'ins167-tdTomato 2aa linkers'. The top 10 fusions, based on brightness and the 593-to-533 ratio, are labeled. The raw data is indicated in Supplementary Table 4.

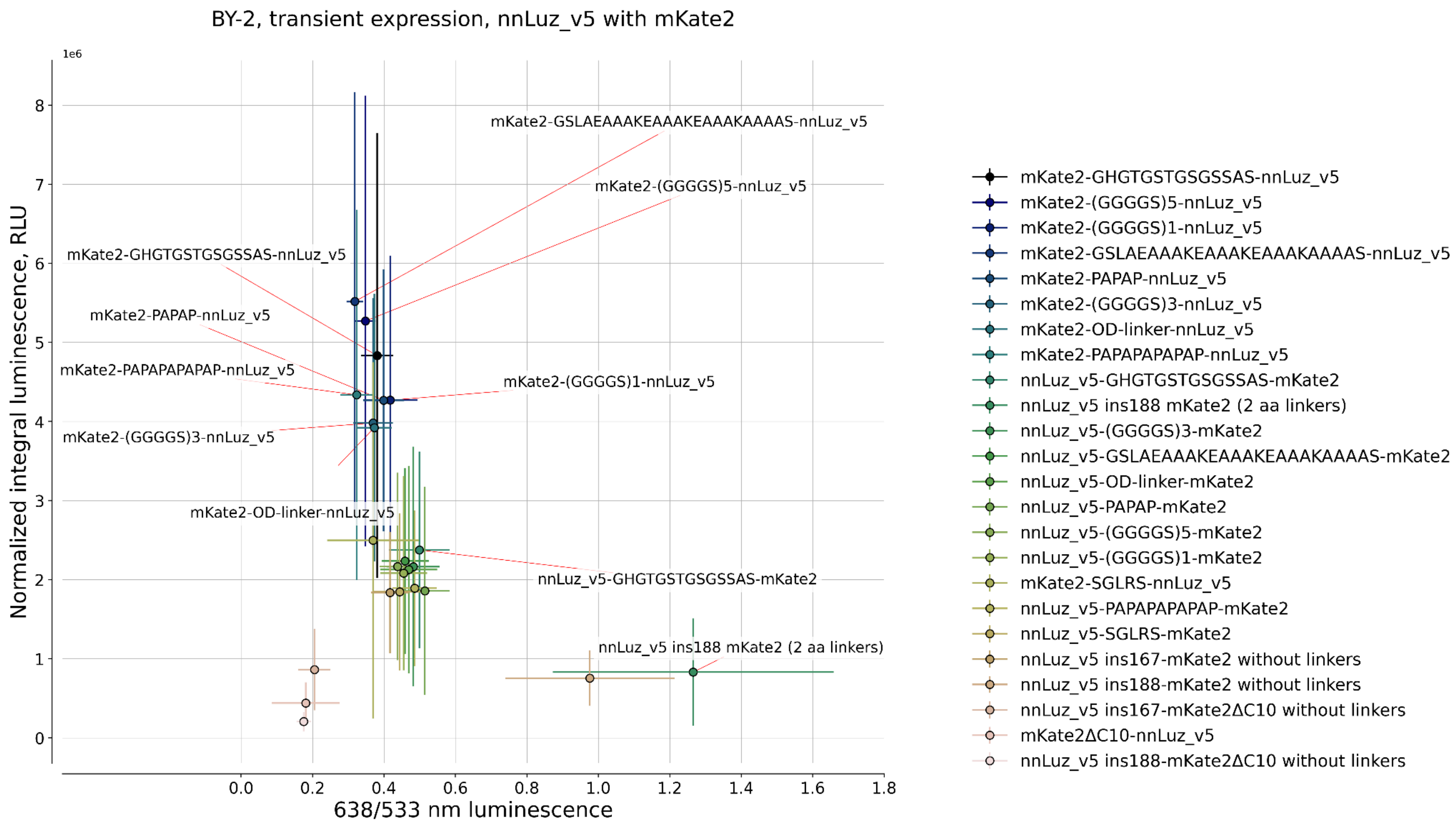

BY-2, transient expression, nnLuz_v5 with mKate2
1e6
Normalized integral luminescence, RLU
638/533 nm luminescence
0.0
0.2
0.4
0.6
0.8
1.0
1.2
1.4
1.6
1.8
0
1
2
3
4
5
6
7
8
mKate2-GSLAEAAAKEAAAKEAAAKAAAAS-nnLuz_v5
mKate2-(GGGGS)5-nnLuz_v5
mKate2-GHGTGSTGSGSSAS-nnLuz_v5
mKate2-PAPAP-nnLuz_v5
mKate2-PAPAPAPAPAP-nnLuz_v5
mKate2-(GGGGS)1-nnLuz_v5
mKate2-(GGGGS)3-nnLuz_v5
mKate2-OD-linker-nnLuz_v5
nnLuz_v5-GHGTGSTGSGSSAS-mKate2
nnLuz_v5 ins188 mKate2 (2 aa linkers)
mKate2-GHGTGSTGSGSSAS-nnLuz_v5
mKate2-(GGGGS)5-nnLuz_v5
mKate2-(GGGGS)1-nnLuz_v5
mKate2-GSLAEAAAKEAAAKEAAAKAAAAS-nnLuz_v5
mKate2-PAPAP-nnLuz_v5
mKate2-(GGGGS)3-nnLuz_v5
mKate2-OD-linker-nnLuz_v5
mKate2-PAPAPAPAPAP-nnLuz_v5
nnLuz_v5-GHGTGSTGSGSSAS-mKate2
nnLuz_v5 ins188 mKate2 (2 aa linkers)
nnLuz_v5-(GGGGS)3-mKate2
nnLuz_v5-GSLAEAAAKEAAAKEAAAKAAAAS-mKate2
nnLuz_v5-OD-linker-mKate2
nnLuz_v5-PAPAP-mKate2
nnLuz_v5-(GGGGS)5-mKate2
nnLuz_v5-(GGGGS)1-mKate2
mKate2-SGLRS-nnLuz_v5
nnLuz_v5-PAPAPAPAPAP-mKate2
nnLuz_v5-SGLRS-mKate2
nnLuz_v5 ins167-mKate2 without linkers
nnLuz_v5 ins188-mKate2 without linkers
nnLuz_v5 ins167-mKate2ΔC10 without linkers
mKate2ΔC10-nnLuz_v5
nnLuz_v5 ins188-mKate2ΔC10 without linkers

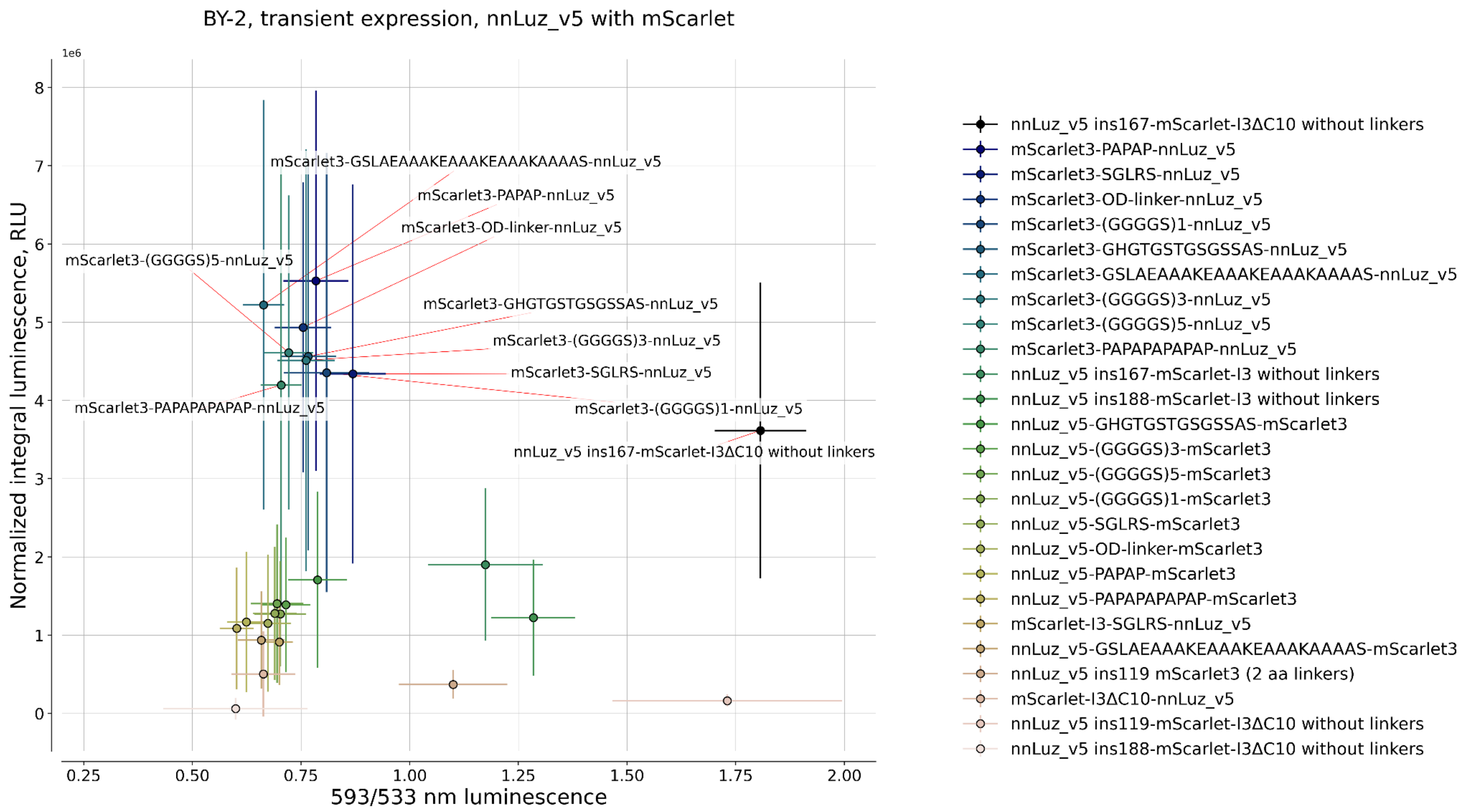
BY-2, transient expression, nnLuz_v5 with mScarlet
1e6
Normalized integral luminescence, RLU
593/533 nm luminescence
0.25
0.50
0.75
1.00
1.25
1.50
1.75
2.00
0
1
2
3
4
5
6
7
8
mScarlet3-GSLAEAAAKEAAAKEAAAKAAAAS-nnLuz_v5
mScarlet3-PAPAP-nnLuz_v5
mScarlet3-OD-linker-nnLuz_v5
mScarlet3-(GGGGS)5-nnLuz_v5
mScarlet3-GHGTGSTGSGSSAS-nnLuz_v5
mScarlet3-(GGGGS)3-nnLuz_v5
mScarlet3-SGLRS-nnLuz_v5
mScarlet3-PAPAPAPAPAP-nnLuz_v5
mScarlet3-(GGGGS)1-nnLuz_v5
nnLuz_v5 ins167-mScarlet-I3ΔC10 without linkers
nnLuz_v5 ins167-mScarlet-I3ΔC10 without linkers
mScarlet3-PAPAP-nnLuz_v5
mScarlet3-SGLRS-nnLuz_v5
mScarlet3-OD-linker-nnLuz_v5
mScarlet3-(GGGGS)1-nnLuz_v5
mScarlet3-GHGTGSTGSGSSAS-nnLuz_v5
mScarlet3-GSLAEAAAKEAAAKEAAAKAAAAS-nnLuz_v5
mScarlet3-(GGGGS)3-nnLuz_v5
mScarlet3-(GGGGS)5-nnLuz_v5
mScarlet3-PAPAPAPAPAP-nnLuz_v5
nnLuz_v5 ins167-mScarlet-I3 without linkers
nnLuz_v5 ins188-mScarlet-I3 without linkers
nnLuz_v5-GHGTGSTGSGSSAS-mScarlet3
nnLuz_v5-(GGGGS)3-mScarlet3
nnLuz_v5-(GGGGS)5-mScarlet3
nnLuz_v5-(GGGGS)1-mScarlet3
nnLuz_v5-SGLRS-mScarlet3
nnLuz_v5-OD-linker-mScarlet3
nnLuz_v5-PAPAP-mScarlet3
nnLuz_v5-PAPAPAPAPAP-mScarlet3
mScarlet-I3-SGLRS-nnLuz_v5
nnLuz_v5-GSLAEAAAKEAAAKEAAAKAAAAS-mScarlet3
nnLuz_v5 ins119 mScarlet3 (2 aa linkers)
mScarlet-I3ΔC10-nnLuz_v5
nnLuz_v5 ins119-mScarlet-I3ΔC10 without linkers
nnLuz_v5 ins188-mScarlet-I3ΔC10 without linkers

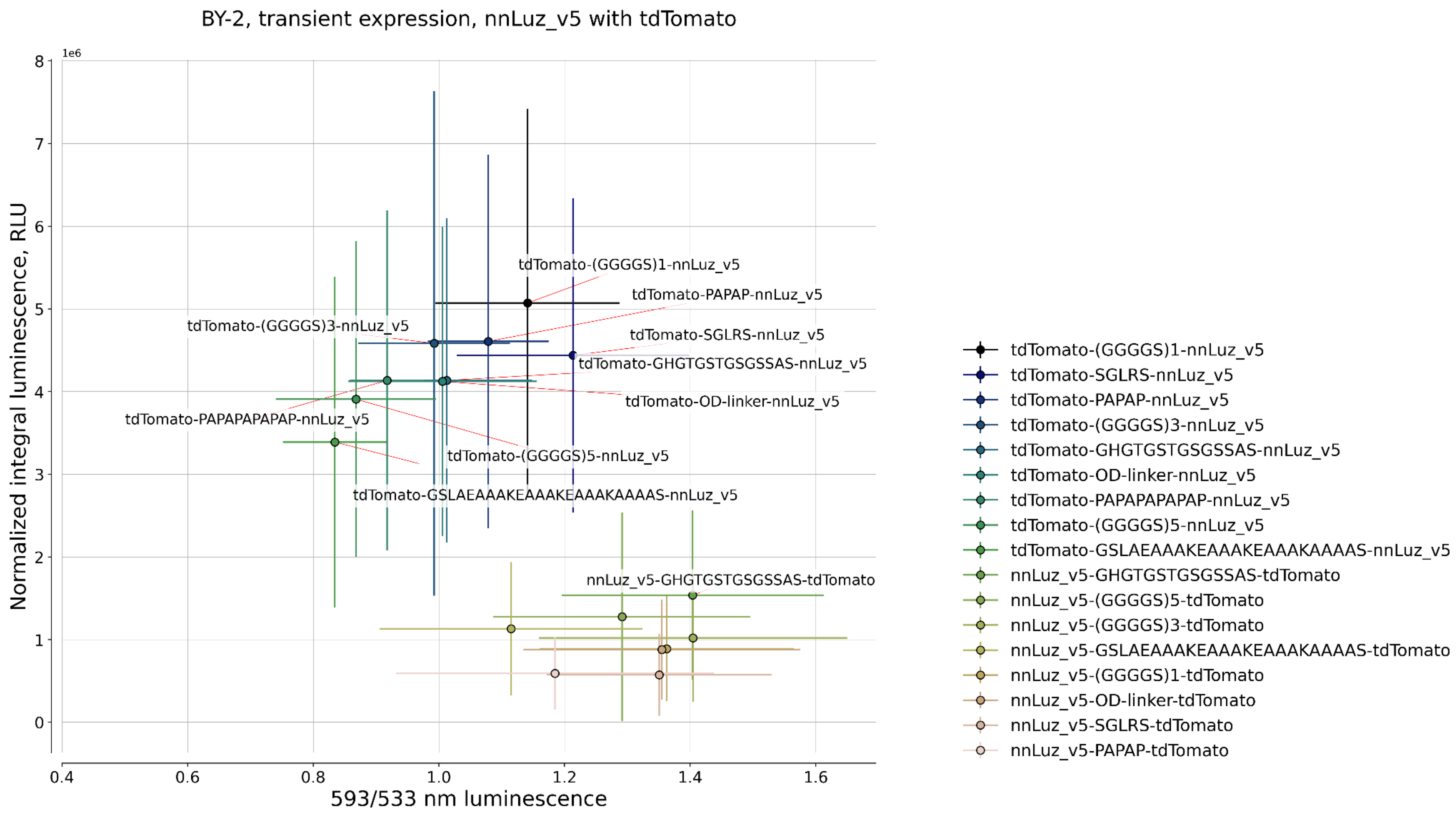


**Supplementary Figure 6. Optimization of bioluminescence resonance energy transfer in nnLuz_v5 through fusion with fluorescent proteins at either the C- or N-terminus of the luciferase.** The top 10 best-performing pairs for each fluorescent protein are labeled. The raw data is indicated in Supplementary Table 5.

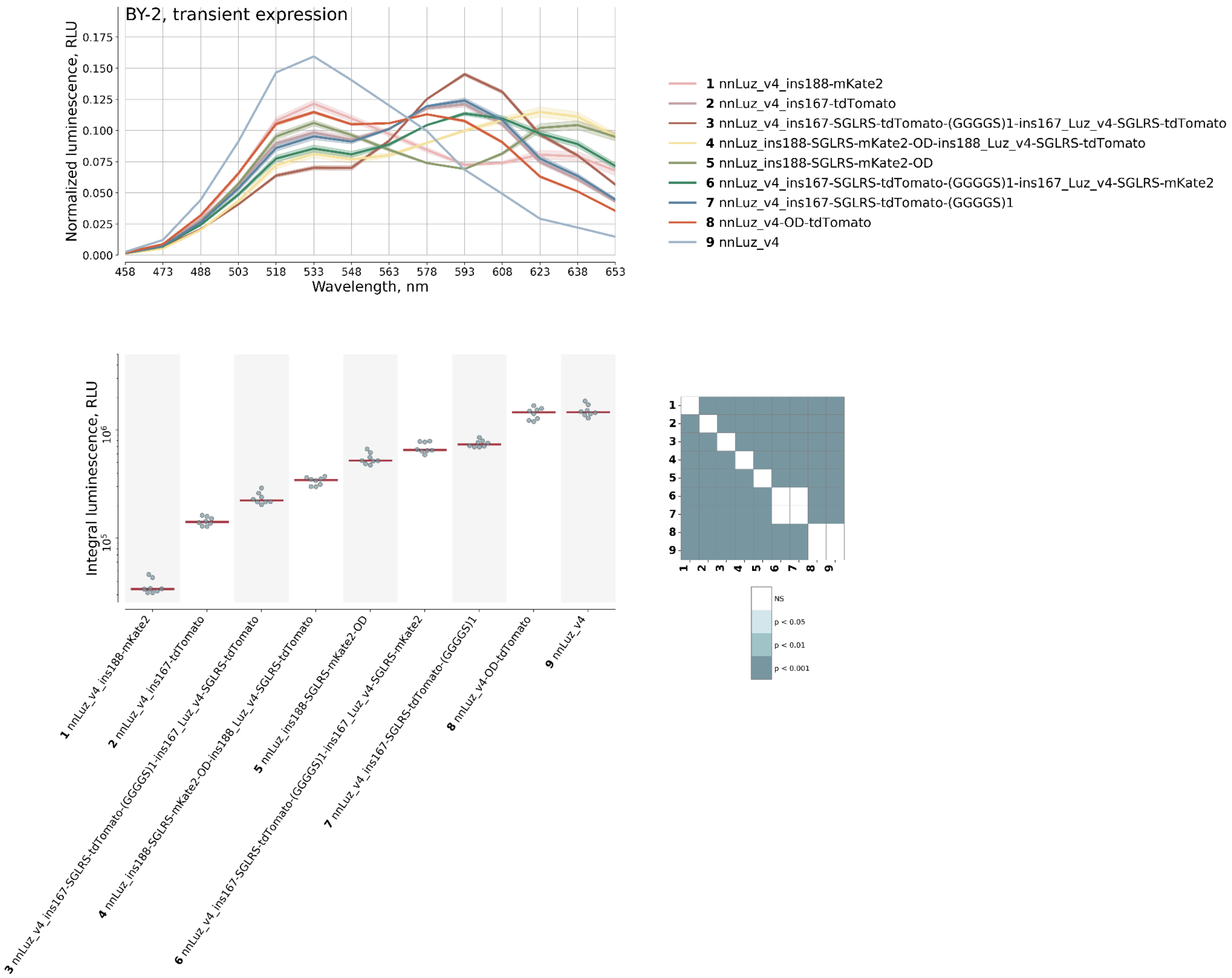


**Supplementary Figure 7. Optimization of bioluminescence resonance energy transfer in already existing BRET pairs through fusion with additional copy of fluorescent proteins at C-terminus of the luciferase.** Spectral characteristics of the fusion proteins are shown at the top; brightness is shown at the bottom.

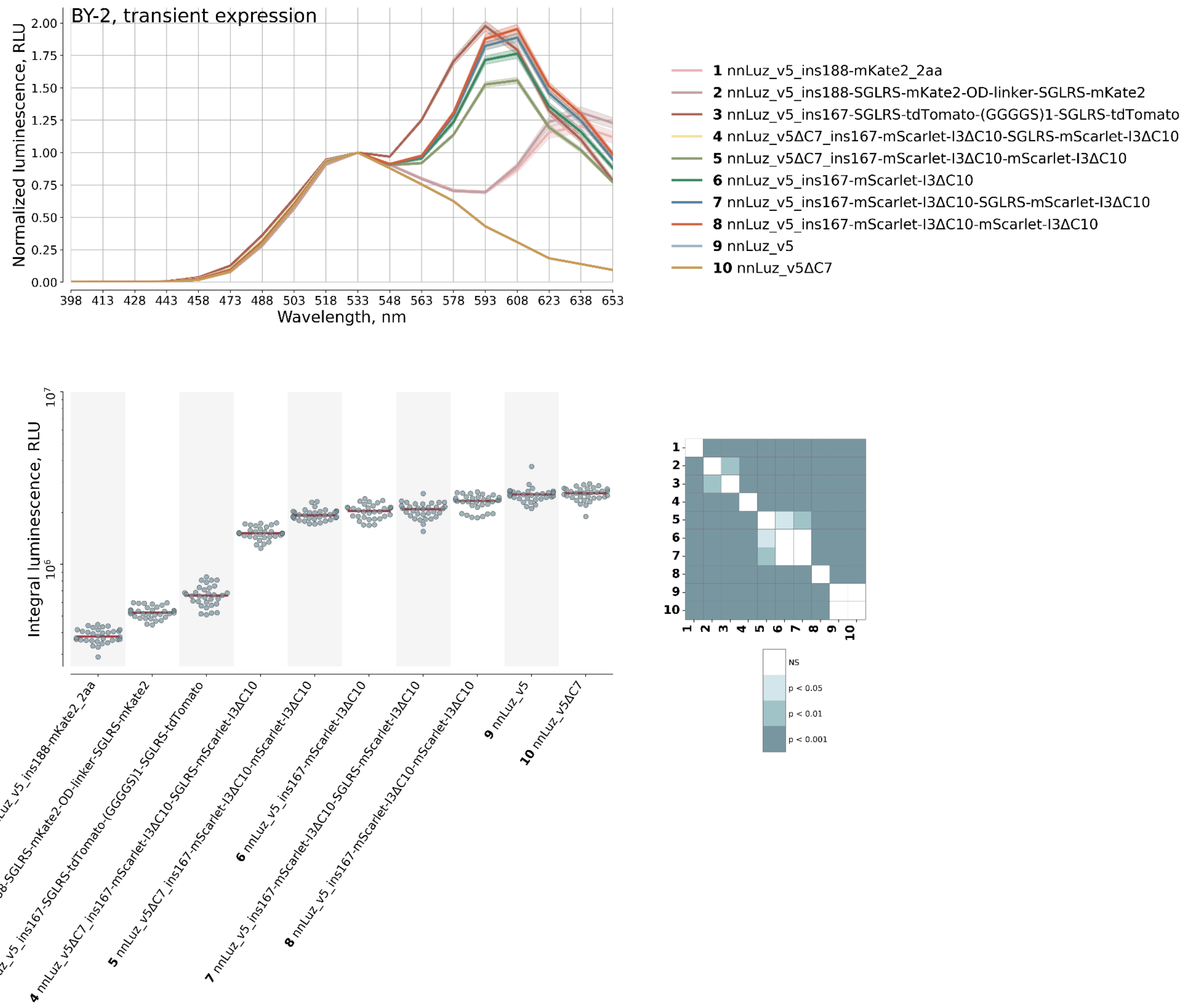


**Supplementary Figure 8. Optimization of bioluminescence resonance energy transfer in already existing BRET pairs through fusion with an additional copy of mScarlet3ΔC10 at C-terminus of the luciferase.** Spectral characteristics of the fusion proteins are shown at the top; brightness is shown at the bottom.

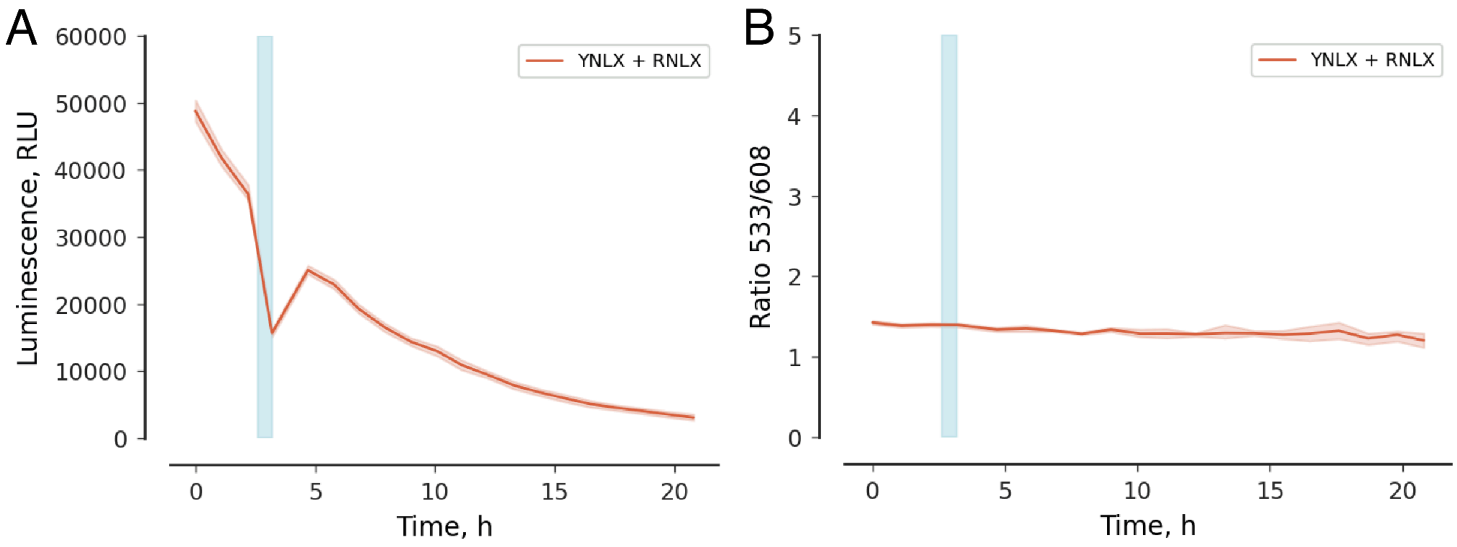


**Supplementary Figure 9. Stability of wavelength ratios in the bacterial luminescence system despite fluctuations in absolute brightness.** Effect of a 30-minute heat shock at 37°C (indicated as a light blue rectangle) on luminescence intensity (A) and ratio 533/608 (B) of luminescence emitted by BY-2 cell packs, expressing yellow (YNLX = VenusΔC10-EL-iluxA), and red (RNLX = mScarlet-IΔC10-EL-iluxA) variants of iluxA in the cytosol, along with the rest of enzymes of bacterial luminescence pathway.

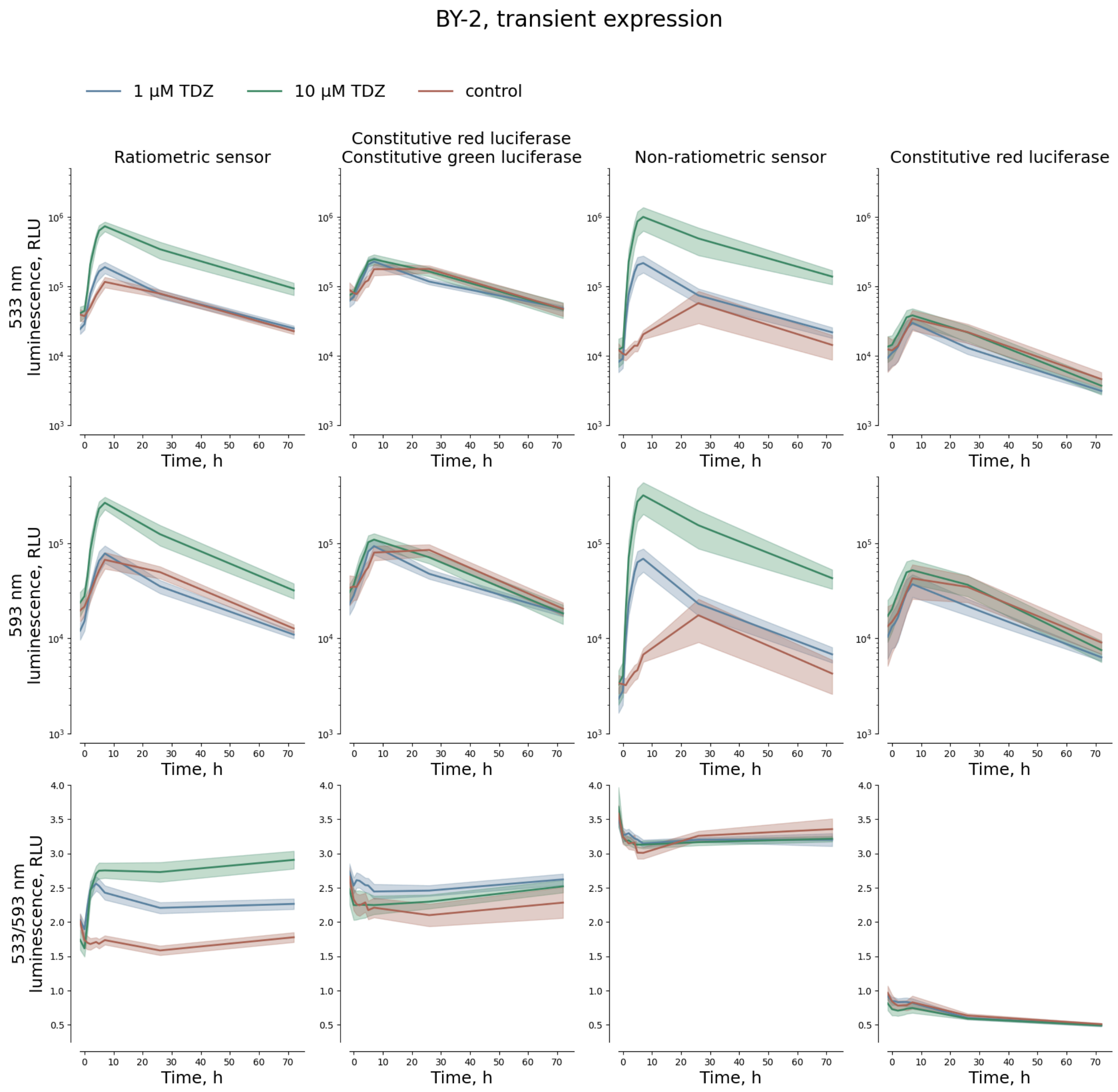


**Supplementary Figure 10. Effect of thidiazuron (TDZ) treatment on the transient expression of fungal ratiometric and non-ratiometric luminescence reporters in *Nicotiana tabacum* BY-2 cell packs.**

Time-course luminescence data showing expression of Luz533 and Luz593 variants driven by different promoter combinations: pTCSv2-Luz533 + MinSyn108-Luz593 (ratiometric sensor), AtTCTP-Luz533 + MinSyn108-Luz593 (constitutive red luciferase + constitutive green luciferase), pTCSv2-Luz533 (non-ratiometric sensor), and MinSyn108-Luz593 (constitutive red luciferase)(columns, left to right). Rows represent measurements of luminescence at 533 nm (top), 593 nm (middle), and the 533/593 nm emission ratio (bottom).

Cell packs were infiltrated with either 1 µM TDZ (blue), 10 µM TDZ (green), or control buffer (brown), and luminescence was monitored over a 72-hour period. Absolute luminescence (RLU) shows TDZ concentration-dependent induction in constructs containing the pTCS2 promoter. In contrast, the 533/593 nm emission ratio remains relatively stable across treatments and time points, suggesting its potential as a ratiometric normalization method.

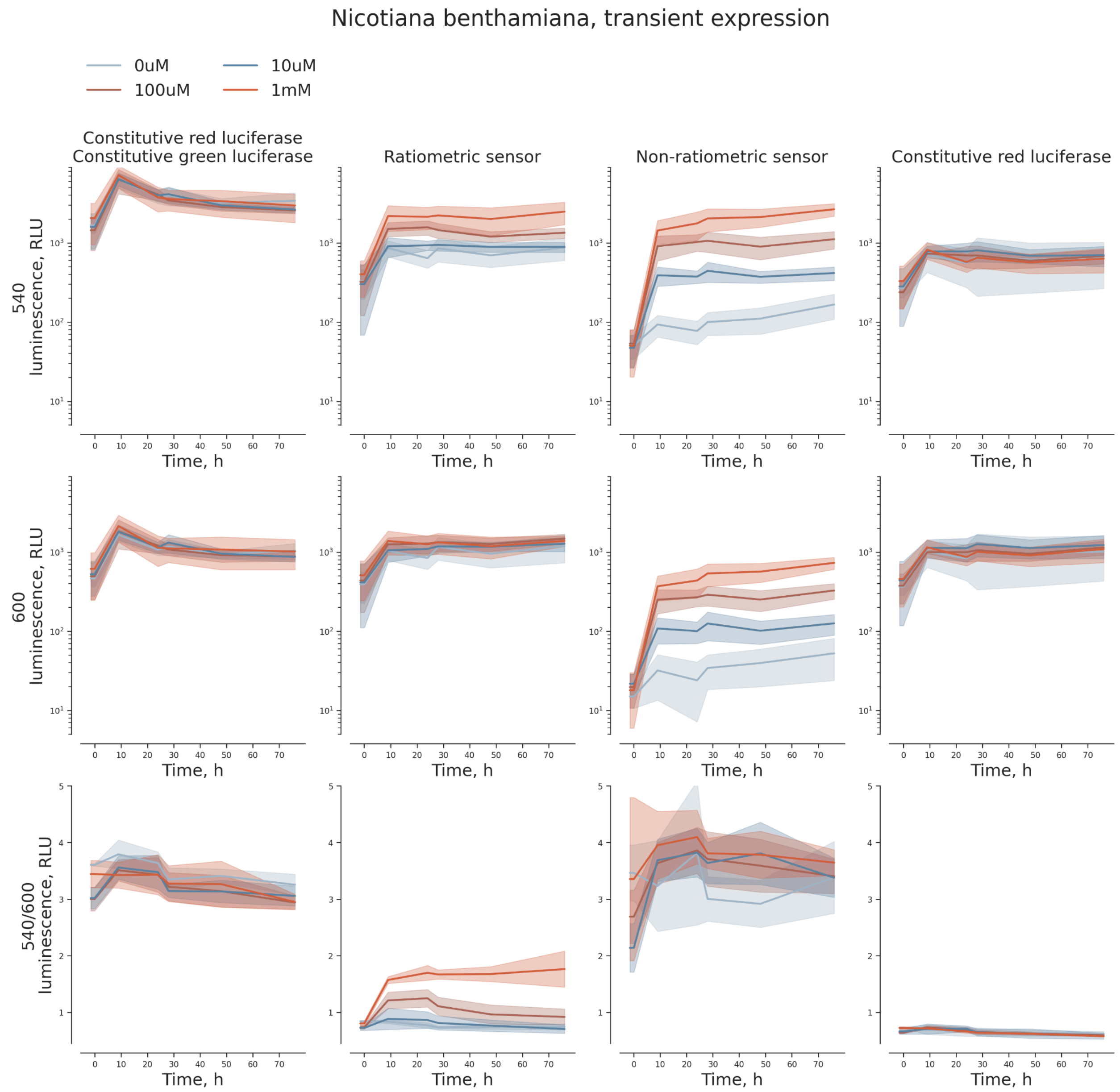


**Supplementary Figure 11. Effect of thidiazuron (TDZ) treatment on the transient expression of fungal ratiometric and non-ratiometric luminescence reporters in *Nicotiana benthamiana* leaves.** Time-course luminescence data showing expression of Luz533 and Luz608 variants driven by different promoter combinations (columns left to right): pTCSv2-Luz533 + p35S-Luz608 (indicated as "Ratiometric sensor"), p35S-Luz533 + p35S-Luz608 (indicated as "Constitutive red luciferase, constitutive green luciferase"), pTCSv2-Luz533 (indicated as "Non-ratiometric sensor"), and p35S-Luz608 (indicated as "Constitutive red luciferase". Rows represent measurements of luminescence at 540 nm (top), 600 nm (middle), and the 540/600 nm emission ratio (bottom). The leaves were  infiltrated with either 1 mM TDZ (light red), 100 μM (brown), 10 μM TDZ (blue), or control buffer (light blue), and luminescence was monitored over a 76-hour period. Absolute luminescence (RLU) shows TDZ concentration-dependent induction in constructs containing the pTCS2 promoter. In contrast, the 540/600 nm emission ratio remains relatively stable across treatments and time points, suggesting its potential as a ratiometric normalization method.

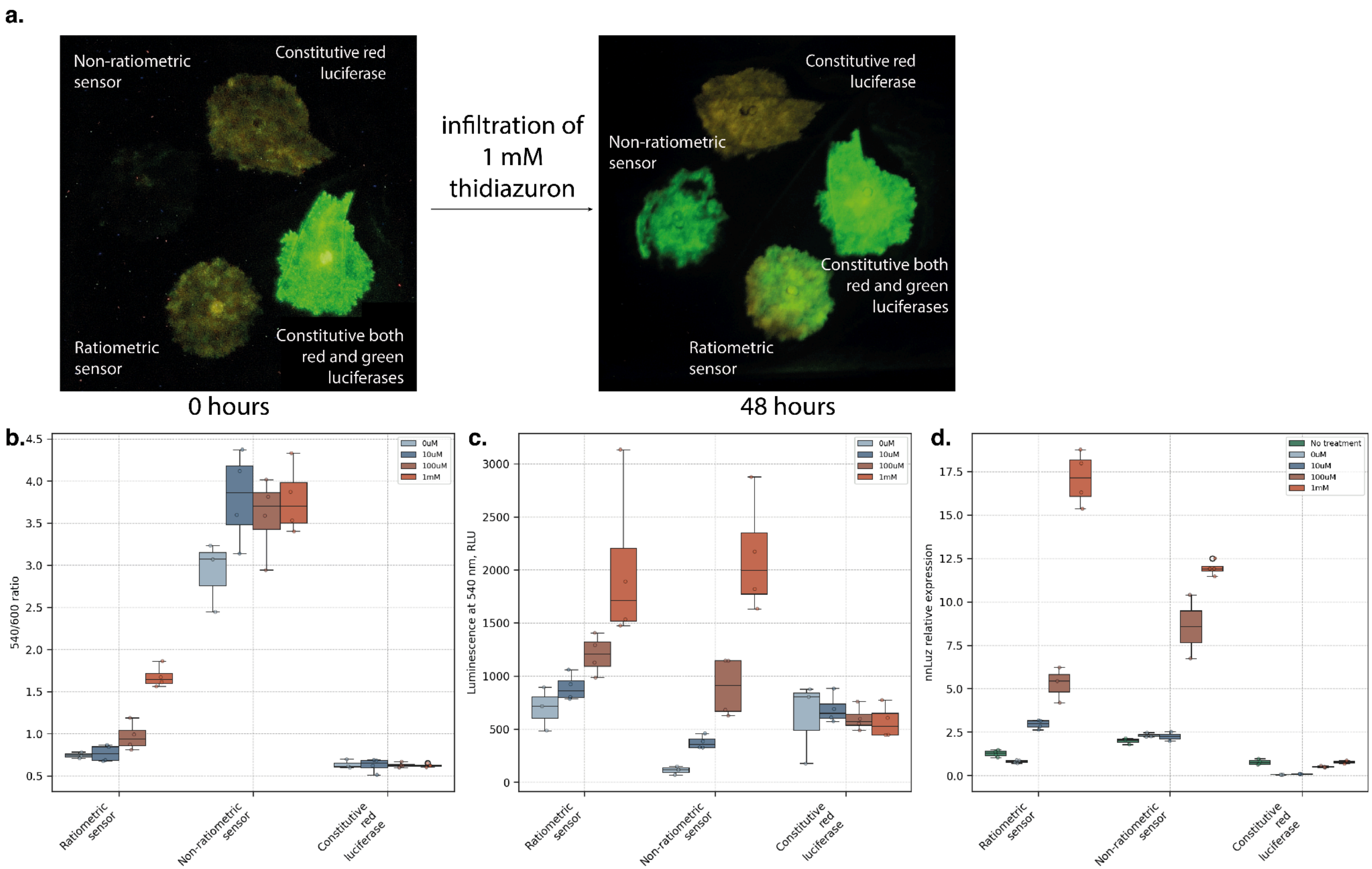


Supplementary Figure 12. Induction of ratiometric (pTCS2-Luz533 + p35S-Luz608), non-ratiometric (pTCS2-Luz533) cytokinin sensors, and controls (constitutive red luciferase – p35S-Luz608; constitutive red and green luciferases – p35S-Luz608 + p35S-Luz533) in *Nicotiana benthamiana* leaves by infiltration with 1 mM thidiazuron solution. a. Representative image of leaves captured using a Sony ILCE-7m3 camera (luminescent signals, ISO 3200, 30-second exposure) before infiltration (‘0 hours’) and 48 hours later ('48 hours'). **b.** Ratio of luminescence values at the reporting wavelength (540 nm) to values at the normalizing wavelength (600 nm). **c.** Absolute luminescence at the reporting wavelength (540 nm). **d.** Luciferase expression levels relative to the housekeeping EF1a gene for the ratiometric and non-ratiometric sensors and the constitutive control.

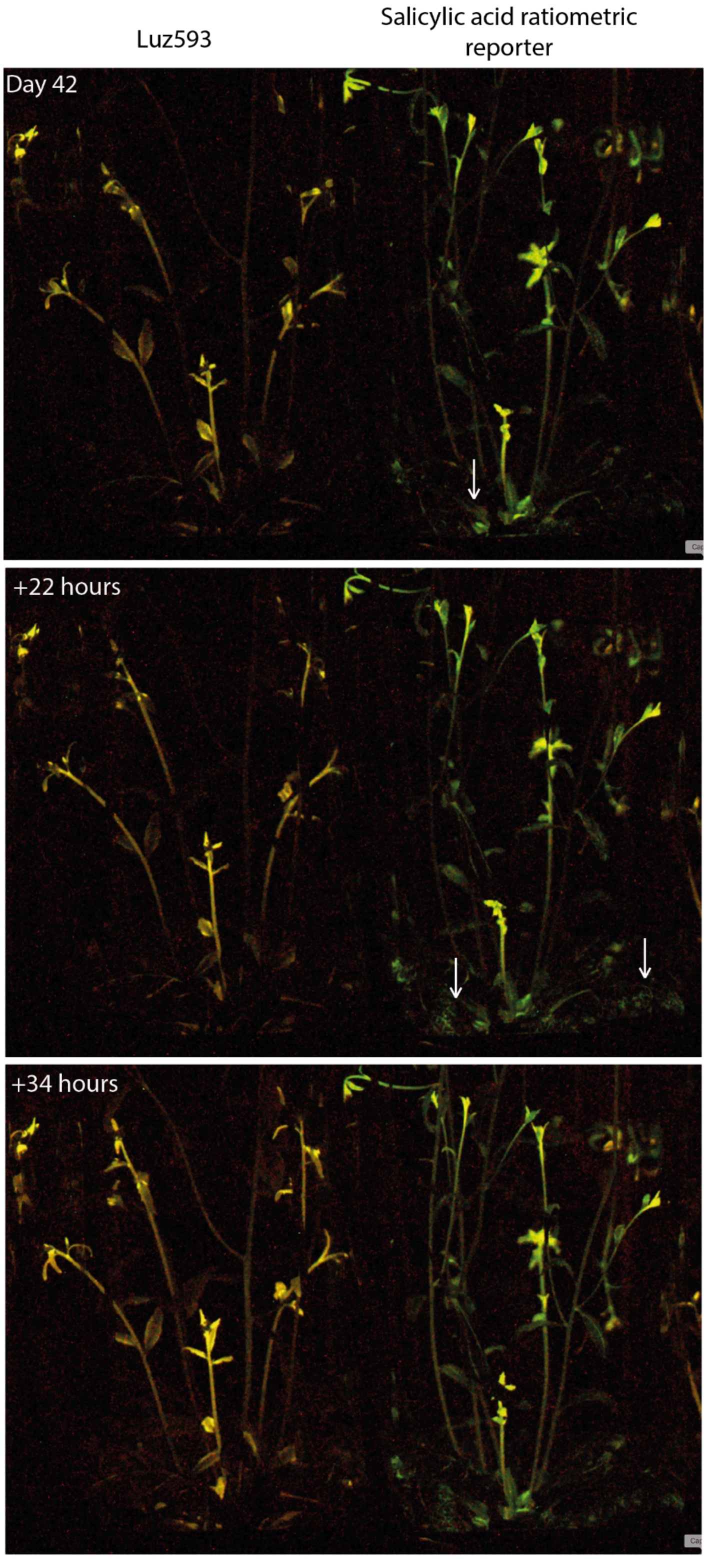


**Supplementary Figure 13. Life-long timelapse snapshots of *Arabidopsis thaliana* plants stably expressing Luz593 or salicylic acid reporter.** Arrows point to dynamic patterns in the leaves during normal plant development.

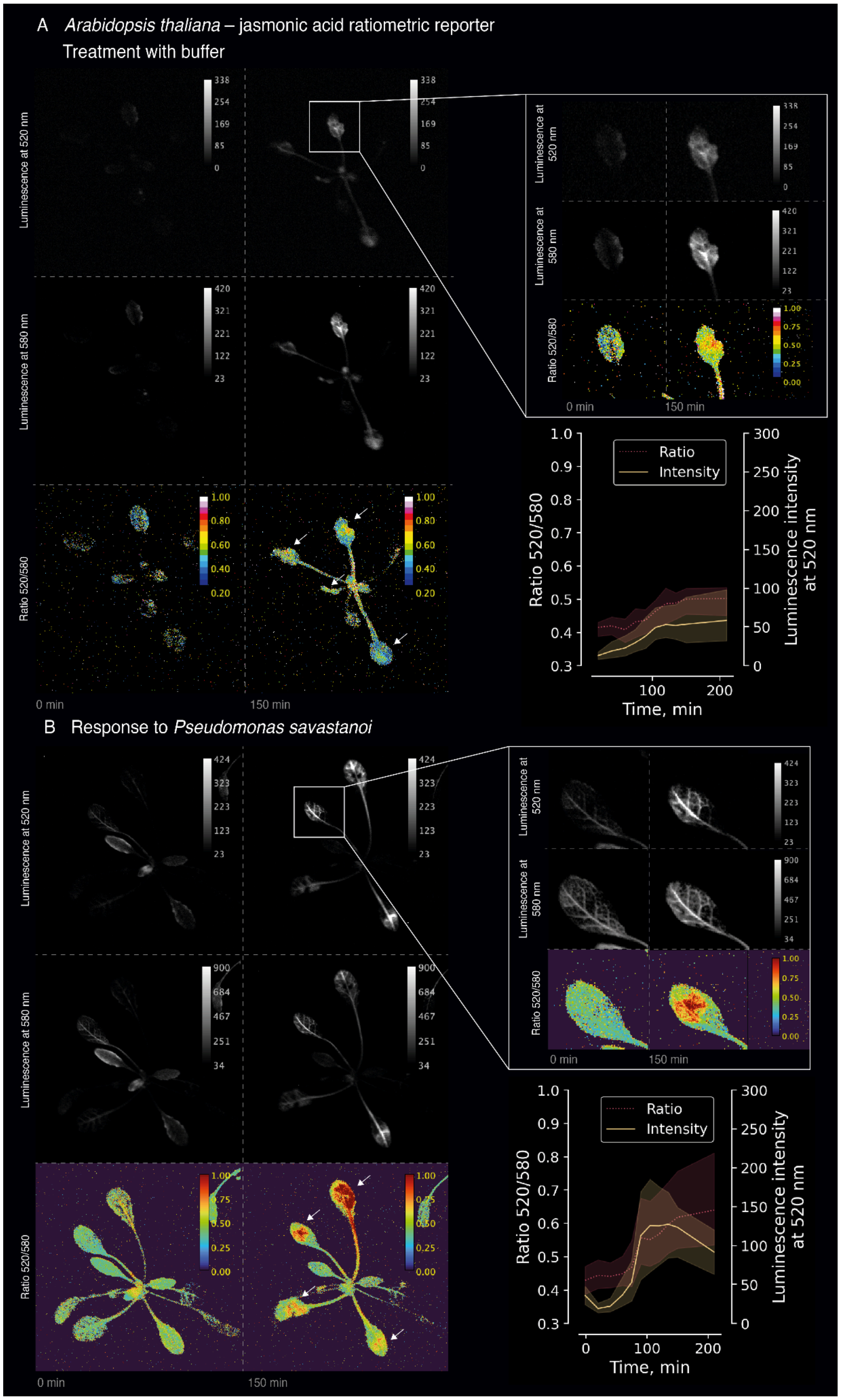


Supplementary Figure 14. Characterization of luminescence ratio and intensity in transgenic *Arabidopsis thaliana* lines expressing jasmonic acid ratiometric reporter treated with buffer (top) or *Pseudomonas savastanoi* (bottom). Red dotted lines indicate the ratio of signals at 520 nm to 580 nm, while yellow solid lines represent luminescence intensity at 520 nm.

Supplementary Video 1. 3-day-long timelapse of 42-44-day-old *Arabidopsis thaliana* plants. Luz593-expressing plant was used as a control, SA ratiometric reporter – salicylic acid ratiometric reporter.

Supplementary Video 2. 53-hour-long timelapse of *Arabidopsis thaliana* plants expressing SA ratiometric reporter – salicylic acid ratiometric reporter.

Supplementary Video 3. 76-hour-long timelapse of transgenic *Arabidopsis thaliana* plants expressing ratiometric jasmonic acid reporter pORCA3-Luz533, pMinSyn108-Luz593. The plants were infiltrated with buffer.

Supplementary Video 4. 76-hour-long timelapse of transgenic *Arabidopsis thaliana* plants expressing ratiometric jasmonic acid reporter pORCA3-Luz533, pMinSyn108-Luz593. The plants were infiltrated with *Pseudomonas savastanoi*.

**Supplementary Table 1.** Plasmids used in the study.

| ID | Plasmid description | Plasmid map |
|---|---|---|
| Y16990 | p35S-Luz608-OCSt | https://benchling.com/s/seq-F6KSPNaZlWDatU6VUjvO?m=slm-zhdzzyjZUa3unN7LrtVm |
| Y17788 | p35S-Luz533-OCSt | https://benchling.com/s/seq-XUzVeTri5PJCwiYkDJzY?m=slm-SdUd3mkfc9oOfz04AHjs |
| Y17790 | p35S-Luz593-OCSt | https://benchling.com/s/seq-HINEE7tvOOIqKjvVDP8L?m=slm-NKE7bCikkLjNs0egjj4E |
| Y17791 | p35S-Luz640-OCSt | https://benchling.com/s/seq-ayr1kOZhAFReGwcbtntb?m=slm-FXLHbSUhsiVcSwDOFNUR |
| Y13213 | pTCS2-min35S-TMV-Luz533-act2t | https://benchling.com/s/seq-QZj4r48LSaY71kGNJcVQ?m=slm-ghLEspiuMXvRsP0HTS4H |
| Y13080 | pAtTCTP-Luz533-act2t | https://benchling.com/s/seq-Uh5EJBY0F4feUhKiWrrw?m=slm-ZRZKmnzNklxzz04esDvi |
| Y9646 | MinSyn108-5'UTR_AtRBCS2B-Luz593-act2t | https://benchling.com/s/seq-daqAR79rc2hZV77lBUit?m=slm-v5S6UrUeaEliyYCG89Mc |
| Y17463 | p35s-nnHispS-OCSt \| pCmYLCV-npgA-ATPt \| dummy \| p35s-nnCPH-OCSt \| pFMV-H3H-NOSt | https://benchling.com/s/seq-JChGOcCVNiYRKbruZQ2z?m=slm-cN2aKkBzOeeBNZLMgSwe |
| dna5168 | p35s-iluxA-OCSt | https://benchling.com/s/seq-3J5UBiU9t6qMrphMhzLF?m=slm-KlSCBeu8PcaiSB3acgN4 |
| dna5169 | p35s-iluxB-OCSt | https://benchling.com/s/seq-I012dBJ0GTrutMoRZHcX?m=slm-Dtc7JE3w6H4BbRfRgymA |
| dna5170 | p35s-iluxC-OCSt | https://benchling.com/s/seq-XfMhkPSbxkCjkPPPWmDx?m=slm-bH4ZjOjLc7EsP6PcMovt |
| dna5171 | p35s-iluxD-OCSt | https://benchling.com/s/seq-T0SKpZl8wDVZdmEy4YbM?m=slm-SDlctnvae3Ceh7aER2zT |
| dna5172 | p35s-iluxE-OCSt | https://benchling.com/s/seq-E5d36yluhQLmwU33eC6y?m=slm-e7hGIht0iHXRNF0QuZry |
| dna5173 | p35s-ilux_frp-OCSt | https://benchling.com/s/seq-9k498FwHyNqd9jG4CRyw?m=slm-YAt9HDKsQAQUbZFvTBJM |
| dna5174 | p35s-VenusΔC10-EL-iluxA-OCSt | https://benchling.com/s/seq-JhbPVfNAnCJfWjHW6it6?m=slm-s97nsWAEjmBVf0U22sc6 |
| dna5175 | p35s-mScarlet-IΔC10-EL-iluxA-OCSt | https://benchling.com/s/seq-yKbsmXHTlZHMz8J35uS2?m=slm-FgzGSjwq9s4J4TCzXpaP |

| | | |
|---|---|---|
| dna5177 | p35s-sfGFPΔC10-EL-iluxA-OCSt | https://benchling.com/s/seq-KUwh8yW8sxDFlw1py8Xn?m=slm-DKQjFkiYJah3Y8BpHodR |

**Supplementary Table 2.** Raw data for Supplementary figure 3.

**Supplementary Table 3.** Raw data for Supplementary figure 4.

**Supplementary Table 4.** Raw data for Supplementary figure 5.

**Supplementary Table 5.** Raw data for Supplementary figure 6.

**Supplementary Table 6.** Plant lines used in the study.

| ID | Plasmid description | Seed generation |
|---|---|---|
| AT7805 | Level M [LM-1] \| [1f] pWRKY70 - nnLuz_v5 - Stop codons - WRKY70_T \| [2r] MinSyn108-5'UTR AtRBCS2B - nnLuz_ins167_SGLRS-tdTomato-(GGGGS)1- SGLRS - tdTomato - act2T \| [4f] HygR cassette | T1 |
| AT7806 | Level M [LM-1] \| [1f] pORCA3 -nnLuz_v5 - Stops codons - ORCA3_T \| [4f] HygR | T1 |
| AT7807 | Level M [LM-1] \| [2r] MinSyn108-5'UTR - nnLuz_ins167_SGLRS-tdTomato-(GGGGS)1- SGLRS - tdTomato - act2T \| [4f] HygR cassette | T1 |
| AT7808 | Level M [LM-1] \| [1f] pORCA3 -nnLuz_v5 - Stops codons - ORCA3_T \| [2r] MinSyn108-5'UTR AtRBCS2B - nnLuz_ins167_SGLRS-tdTomato-(GGGGS)1- SGLRS - tdTomato - act2T \| [4f] HygR cassette | T1 |

### Methods

#### Design and assembly of genetic constructs

Coding sequences of bioluminescence genes were optimised for the expression in *Nicotiana benthamiana* and *Arabidopsis thaliana* and ordered synthetically (**Supplementary Table 1**). Golden Gate assembly was performed in the T4 ligase buffer (Thermo Fisher) containing 10 U of T4 ligase, 20 U of either BsaI, BpiI, BtgZI, MlyI or BsmBI (Thermo Fisher) and ~100 ng of each DNA part. Typically, Golden Gate reactions were performed according to 'troubleshooting' cycling conditions described in [ref [1]]: 25 cycles (90 s at 37°C, 180 s at 16°C), then 5 min at 50°C and 10 min at 80°C.

Correct DNA assembly was typically confirmed by Sanger sequencing, and in some cases additionally by Nanopore whole plasmid sequencing. DNA assembly and whole-plasmid sequencing was typically ordered from Cloning Facility (cloning.tech).

#### Transformation of *Arabidopsis thaliana*

*Arabidopsis thaliana* (ecotype Columbia 0) was transformed with floral dip [2] with a binary vector carrying the transcriptional units for the production of the fungal luciferin, and the recycling of the luciferin's oxidization product (caffeoylpyruvic acid). To generate stable *Arabidopsis thaliana* plants expressing either salicylic- or jasmonic-acid-responsive reporters, we used a "luciferase-less" masterline carrying genes necessary for fungal luminescence, except the luciferase gene. This masterline was already generated by Michael Karampelias in our previous study [3]. To obtain such a masterline Level P plasmid pNK091 was used, it carried (1) nnCPH gene under the control of 0.4 kb constitutive 35S promoter from cauliflower mosaic virus with 5' untranslated region of TMV omega virus and ocs terminator from *Agrobacterium tumefaciens*; (2) wild-type version of nnH3H gene under the control of constitutive FMV promoter from figwort mosaic virus and nopaline synthase terminator from *Agrobacterium tumefaciens*; (3) nnHispS gene under the control of 0.4 kb constitutive 35S promoter from cauliflower mosaic virus with 5' untranslated region of TMV omega virus and ocs terminator from *Agrobacterium tumefaciens*; (4) NpgA gene under the control of the constitutive CmYLCV 9.11 promoter from Cestrum yellow leaf curling virus and ATPase terminator from *Solanum lycopersicum*; (5) kanamycin resistance cassette driven by pNos promoter and ocs terminator from *Agrobacterium tumefaciens*.

After four generations of antibiotic resistance (kanamycin), homozygous plants were transformed with plasmids encoding constitutively expressed Luz593, ratiometric jasmonic acid reporter or ratiometric salicylic acid reporter-carrying transcription units. To select the transformants the plasmids carried hygromycin resistance cassette driven by pNos promoter and ocs terminator from *Agrobacterium tumefaciens*.

#### Transformation of *Agrobacterium tumefaciens*

Plasmids were transformed into competent cells of *Agrobacterium tumefaciens* AGL0 [4], and clones were selected on LB (Luria-Bertani) agar plates containing 50 mg/L of rifampicin and an additional antibiotic, depending on the plasmid used for transformation (200 mg/L of carbenicillin, 50 mg/mL of kanamycin or 100 mg/mL spectinomycin). Individual colonies were then inoculated into 10 ml of LB medium containing the same concentration of antibiotics. After overnight incubation at 28°C with shaking at 220 rpm, cultures were centrifuged at 2900 g, resuspended in 25% glycerol and stored as glycerol stocks at −80 °C.

Growth conditions for *Arabidopsis thaliana*

Seeds were surface sterilised with 70% ethanol for 15 minutes and washed twice with 100% ethanol. Seeds were dried in a laminar flow and then sown on the surface of solid (10% plant agar) half-strength MS medium with 1% sucrose. Sown seeds were vernalized for 2 days at 4 °C and then transferred to grow vertically at 22 °C.

Validation of reporters in plant cell packs based on *Nicotiana tabacum* BY-2 cell culture

BY-2 cell culture was grown in BY-2 medium (Murashige and Skoog (MS) with 0.2 mg/L 2,4-dichlorophenoxyacetic acid, 200 mg/L $KH_2PO_4$, 1 mg/L thiamine, 100 mg/L myo-inositol and 30 g/L sucrose) at 27°C by shaking at 130 rpm in darkness, with 2 ml of one-week-old culture being transferred into new 200 ml of BY-2 medium every week [5].

Transformations of BY-2 cell packs were made according to a protocol adapted from [ref. [6]]. One-week-old BY-2 culture was pelleted in black 96-well plates to create cell packs that were infiltrated by a mixture of several agrobacterial strains containing binary vectors. One of the strains encoded silencing inhibitor P19 ($OD_{600}$0.2), and other encoded bioluminescence genes and a reporter (pTSCv2-driven ratiometric or non-ratiometric variants) gene ($OD_{600}$0.5). Comparison of reporters was done by co-infiltrating BY-2 cell packs with agrobacterium individually encoding bioluminescent enzymes.

Cytokinin reporter functionality was tested in BY-2 cell packs. The cell packs were treated with 1 µM, 10 µM or mock  (0 µM) thidiazuron (TDZ) solution. TDZ was dissolved in an infiltration buffer. The 100 µl of TDZ solutions was applied on the cell packs, the excess of the solution was removed with centrifugation for 1 min, 500 g. Signal variation was evaluated by calculating the ratio of mean luminescence intensity at the peak (7 hours post-treatment) to that at the end of the experiment (72 hours post-treatment).

For elucidation of wavelength ratio stability for both fungal and bacterial bioluminescence color variants the following procedures were used: (1) treatment with infiltration buffer [7], (2) treatment with 200 mg/L of quizalofop-P-ethyl herbicide, (3) heat shock at 37 °C for 30 minutes. The 100 µL of the solutions was applied on the cell packs, the excess of the liquid was removed with centrifugation for 1 min, 500g.

After the treatments the plates were incubated at 80% humidity and 22 °C and immediately imaged for 48 hours using consumer-grade Sony camera (see details below).

Validation of reporters in *Nicotiana benthamiana* leaves

Four-week-old wild-type *N. benthamiana* plants (NB000) were infiltrated with the mixtures of agrobacteria *A. tumefaciens* AGL0 strains expressing p19 (OD 0.1), fungal luciferin biosynthesis and recycling genes, and the following compositions: (1) p35S-Luz533 and p35S-Luz608, (2) pTCSv2-Luz533 and p35S-Luz608, (3) p35S-Luz608, (4) pTCSv2-Luz533. All the sample infiltration spots were located at the same leaves.

Seventy two hours post initial infiltrations the same spots were additionally infiltrated with thidiazuron solutions 0 µM, 10 µM, 100 µM, 1 mM. At several time points up to 76 hours post infiltration, photos were captured with Sony camera and IVIS imager at luminescence emission wavelengths 500-700 nm.

For qPCR assays housekeeping gene EF1a was used with primers: EF1a-qrtF CTGCAACAAGATGGATGCTAC, EF1a-qrtR ACCAACAGGGACAGTACCAAT [8]. To assess nnLuz expression level the following primers were used: nnLuz_v5-qrtF CTCCCACATTATACAGCGTCAGCGT, nnLuz_v5-qrtR TGTGTGTCTTGCTTGCACACG. For amplification the following parameters were utilised: (preincubation: 95 °C — 3 min; 3-step amplification (40 cycles): 95 °C — 15 sec, 60 °C — 20 sec, 72 °C — 30 sec; cooling: 20 °C — 1 hour).

##### Validation of phytohormone reporters in transgenic *Arabidopsis thaliana*

We infected transgenic plants with hemibiotrophic pathogen *Pseudomonas savastanoi* (a pathovar of *Pseudomonas syringae)*. To infect the plants we infiltrated the leaves with the bacteria solution with the $OD_{600}$ of bacteria was 0.4. The bacteria were resuspended in an infiltration buffer described in [ref. [7]]. The buffer was used as a mock control.

##### Luminescent imaging on consumer-grade camera

The plants were imaged for 48 hours using Sony Alpha ILCE-7M3 camera and 35-mm T1.5 ED AS UMC VDSLR lens (Samyang, ~f/1.4) with an exposure time of 5-30 s and ISO values 400, 3,200, and 20,000. The plants typically were imaged every 30 minutes. Processing of images was performed using FiJi ImageJ distribution (version 1.53t) and custom Python scripts. Background-subtracted mean values in the region of interest were used for luminescence quantification. Background subtraction was performed using the following formula:
$signal = signal_{raw} - background_{mean}$.

##### Plant imaging on IVIS Spectrum CT

The imaging of *Pseudomonas savastanoi* infection was performed using IVIS Spectrum CT, with filter for 520 and 580 nm in front of the camera, an exposure time of 30 s, and binning 2. The samples were acquired with “D” field of view settings. Ambient light image was taken after the luminescence measurements. Other settings were left at their default values.

##### Image analysis

All images were analyzed and processed with Fiji (ImageJ, NIH). Images were converted to 32 bits and a 580-nm stack was thresholded to eliminate pixel values from background (Not-a-Number function). The 520-nm stack was divided into the 580-nm stack frame-by-frame.

The resulting stack was depicted in pseudocolors using a “Turbo” lookup table. The ratio was calculated for the region of interest (ROI). The resulting ratio data were subsequently analyzed with custom Python scripts.

##### Data presentation and statistical analysis

Most of the data were plotted as medians and coloured individual data points using Seaborn (https://seaborn.pydata.org/, ver. 0.13.2) and Matplotlib (https://matplotlib.org/, ver. 3.8.0) packages, using Python version 3.10.12. Pairwise post-hoc two-sided Mann–Whitney U tests (Scikit-posthocs package [9], version 0.10.0) were computed. Sample numbers (N) were reported in the figures.
