## Supplementary Table 2 for "Metabolically robust autoluminescent reporters"

| | Integral luminescence, mean | 640/533 luminescence, mean |
|---|---|---|
| Label on the plot | | |
| SGLRS-(GGGGS)5 | 4732.78365 | 1.180279983 |
| SGLRS-SGLRS | 4960.57315 | 1.123914959 |
| (GGGGS)3-SGLRS | 5531.32415 | 0.9980595535 |
| GHGTGSTGSGSSAS-(GGGGS)1 | 5684.874025 | 0.9646612973 |
| SGLRS-(GGGGS)3 | 4293.43115 | 1.2077278 |
| SGLRS-OD-linker | 4077.094275 | 1.26896714 |
| SGLRS-PAPAPAPAPAP | 4403.613525 | 1.172648555 |
| OD-linker-(GGGGS)1 | 5255.073275 | 0.9755081691 |
| (GGGGS)1-(GGGGS)1 | 4422.3844 | 1.147116924 |
| PAPAP-(GGGGS)1 | 4704.996025 | 1.075498283 |
| SGLRS-(GGGGS)1 | 4187.367775 | 1.203169144 |
| ins167-mKate2 without linkers | 10639.48928 | 0.4172438886 |
| ins188-mKate2 2aa linkers | 3109.428525 | 1.395996133 |
| OD-linker-SGLRS | 4396.016025 | 0.9804733309 |
| (GGGGS)1-SGLRS | 3934.401275 | 1.076561403 |
| (GGGGS)3-(GGGGS)1 | 3999.049525 | 1.041572018 |
| PAPAPAPAPAP-(GGGGS)1 | 4983.35065 | 0.808821368 |
| (GGGGS)5-(GGGGS)1 | 4565.981775 | 0.875410518 |
| SGLRS-PAPAP | 3681.08415 | 1.045934083 |
| PAPAP-SGLRS | 3346.06965 | 1.10549167 |
| GHGTGSTGSGSSAS-PAPAP | 4252.914275 | 0.8548804543 |
| SGLRS-GSLAEAAAKEAAAKEAAAKAAAAS | 3196.61015 | 1.125065762 |
| (GGGGS)3-(GGGGS)5 | 4554.22565 | 0.7821272066 |
| SGLRS-GHGTGSTGSGSSAS | 2923.90115 | 1.212798724 |
| GHGTGSTGSGSSAS-SGLRS | 3498.96465 | 1.008084551 |
| PAPAP-PAPAPAPAPAP | 3791.5734 | 0.916652532 |
| GHGTGSTGSGSSAS-(GGGGS)3 | 3994.62715 | 0.8489209308 |
| (GGGGS)5-(GGGGS)3 | 4506.7589 | 0.7398556747 |
| GSLAEAAAKEAAAKEAAAKAAAAS-(GGGGS)1 | 4915.421525 | 0.6705676666 |
| (GGGGS)3-(GGGGS)3 | 3778.18765 | 0.8613700211 |

| | | |
|---|---|---|
| OD-linker-(GGGGS)3 | 3906.827275 | 0.8213674534 |
| GHGTGSTGSGSSAS-OD-linker | 4040.831025 | 0.791726299 |
| PAPAP-(GGGGS)5 | 3258.0464 | 0.9772734768 |
| PAPAPAPAPAP-OD-linker | 4853.75915 | 0.6512665556 |
| PAPAP-PAPAP | 3253.86065 | 0.9678537244 |
| PAPAP-GHGTGSTGSGSSAS | 3087.98615 | 1.00550653 |
| PAPAP-OD-linker | 3134.1514 | 0.9820010846 |
| PAPAPAPAPAP-PAPAP | 4348.24215 | 0.7017600208 |
| OD-linker-PAPAP | 3630.5254 | 0.8237259549 |
| (GGGGS)5-SGLRS | 3360.43515 | 0.8801489135 |
| PAPAPAPAPAP-SGLRS | 3655.3269 | 0.8089773956 |
| (GGGGS)1-(GGGGS)5 | 2889.6314 | 0.9956884381 |
| OD-linker-OD-linker | 3766.887775 | 0.7542710324 |
| (GGGGS)1-PAPAPAPAPAP | 2985.310025 | 0.935224046 |
| PAPAP-(GGGGS)3 | 2748.468525 | 1.015517087 |
| (GGGGS)1-OD-linker | 2715.033275 | 1.027588483 |
| (GGGGS)1-(GGGGS)3 | 2652.609029 | 1.050892228 |
| GHGTGSTGSGSSAS-GHGTGSTGSGSSAS | 3253.293275 | 0.8384530043 |
| OD-linker-PAPAPAPAPAP | 3976.377525 | 0.6828145969 |
| GHGTGSTGSGSSAS-PAPAPAPAPAP | 3811.314025 | 0.7092298887 |
| (GGGGS)1-PAPAP | 2673.96915 | 0.9903018995 |
| (GGGGS)5-PAPAP | 3454.3574 | 0.748657393 |
| (GGGGS)3-OD-linker | 3145.727275 | 0.8203283796 |
| (GGGGS)3-PAPAP | 2906.934775 | 0.8768698206 |
| OD-linker-(GGGGS)5 | 3351.6449 | 0.7519658117 |
| (GGGGS)1-GHGTGSTGSGSSAS | 2458.15115 | 1.024287857 |
| (GGGGS)5-OD-linker | 3541.52015 | 0.7001949559 |
| (GGGGS)3-PAPAPAPAPAP | 3383.7049 | 0.7225598802 |
| PAPAPAPAPAP-PAPAPAPAPAP | 4323.019025 | 0.5592894916 |
| GSLAEAAAKEAAAKEAAAKAAAAS-SGLRS | 3593.678275 | 0.6665049653 |
| GSLAEAAAKEAAAKEAAAKAAAAS-PAPAP | 3997.114275 | 0.5873674784 |

| | | |
|---|---|---|
| (GGGGS)5-GHGTGSTGSGSSAS | 3203.248275 | 0.7317673602 |
| nnLuz_v5 | 19915.47553 | 0.1173861506 |
| GHGTGSTGSGSSAS-(GGGGS)5 | 3039.756275 | 0.7670304514 |
| OD-linker-GHGTGSTGSGSSAS | 2806.6984 | 0.8141492719 |
| PAPAPAPAPAP-GHGTGSTGSGSSAS | 3301.00265 | 0.6831121286 |
| ins188-mKate2 without linkers | 1707.56065 | 1.13747487 |
| PAPAPAPAPAP-(GGGGS)3 | 2786.2324 | 0.6909831237 |
| PAPAPAPAPAP-(GGGGS)5 | 2990.806525 | 0.6422226217 |
| (GGGGS)5-(GGGGS)5 | 2881.080025 | 0.666151503 |
| GHGTGSTGSGSSAS-GSLAEAAAKEAAAKEAAAKAAAAS | 2983.729525 | 0.6369035834 |
| (GGGGS)5-PAPAPAPAPAP | 3222.644025 | 0.5867757336 |
| PAPAP-GSLAEAAAKEAAAKEAAAKAAAAS | 2272.43165 | 0.8317428757 |
| GSLAEAAAKEAAAKEAAAKAAAAS-OD-linker | 3577.6549 | 0.5159864714 |
| (GGGGS)3-GHGTGSTGSGSSAS | 2095.019025 | 0.8770853263 |
| (GGGGS)1-GSLAEAAAKEAAAKEAAAKAAAAS | 2146.40815 | 0.8543746258 |
| GSLAEAAAKEAAAKEAAAKAAAAS-(GGGGS)3 | 3170.941525 | 0.5679037165 |
| (GGGGS)5-GSLAEAAAKEAAAKEAAAKAAAAS | 3336.9619 | 0.5286135469 |
| OD-linker-GSLAEAAAKEAAAKEAAAKAAAAS | 2963.732275 | 0.588585909 |
| GSLAEAAAKEAAAKEAAAKAAAAS-GHGTGSTGSGSSAS | 3102.51565 | 0.5601424241 |
| GSLAEAAAKEAAAKEAAAKAAAAS-PAPAPAPAPAP | 3722.13015 | 0.4560995066 |
| (GGGGS)3-GSLAEAAAKEAAAKEAAAKAAAAS | 2446.729025 | 0.6438876593 |
| GSLAEAAAKEAAAKEAAAKAAAAS-(GGGGS)5 | 3138.627025 | 0.4915610265 |
| GSLAEAAAKEAAAKEAAAKAAAAS-GSLAEAAAKEAAAKEAAAKAAAAS | 2575.526275 | 0.4092753321 |
| PAPAPAPAPAP-GSLAEAAAKEAAAKEAAAKAAAAS | 2002.796275 | 0.5206195923 |
| ins167-mKate2ΔC10 without linkers | 3679.850525 | 0.1536468422 |
| mKate2-SGLRS-nnLuz_v5 | 995.495525 | 0.1995783823 |
| mKate2ΔC10-nnLuz_v5 | 1214.579775 | 0.1176466198 |
| ins188-mKate2ΔC10 without linkers | 810.6539143 | 0.1324181604 |
| empty | -0.5110583333 | 0.2180720622 |
