## Supplementary Table 3 for "Metabolically robust autoluminescent reporters"

| | Integral luminescence, mean | 608/533 luminescence, mean |
|---|---|---|
| Label on the plot | | |
| ins167-mScarlet-I3ΔC10 without linerks | 1596653.8 | 1.734178758 |
| nnLuz_v5 | 2636097.6 | 0.3038782378 |
| ins167-mScarlet-I3 without linkers | 661251 | 1.098332482 |
| ins188-mScarlet-I3 without linkers | 307417.4 | 1.174968711 |
| (GGGGS)5-(GGGGS)3 | 118015.4 | 1.008205286 |
| mScarlet-I3-SGLRS-nnLuz_v5 | 206583 | 0.5661293534 |
| GHGTGSTGSGSSAS-PAPAP | 99097 | 1.041423218 |
| ins119 mScarlet3 (2 aa linkers) | 92658.2 | 1.107461885 |
| PAPAPAPAPAP-PAPAP | 98292.2 | 1.01888524 |
| PAPAPAPAPAP-OD-linker | 122298.6 | 0.8112395063 |
| (GGGGS)3-OD-linker | 113410.8 | 0.8654906411 |
| GSLAEAAAKEAAAKEAAAKAAAAS-GHGTGSTGSGSSAS | 99184.8 | 0.9693844935 |
| SGLRS-GSLAEAAAKEAAAKEAAAKAAAAS | 104454.4 | 0.8849043125 |
| PAPAP-PAPAP | 89013 | 1.037041088 |
| OD-linker-OD-linker | 98418.8 | 0.9076267258 |
| OD-linker-GHGTGSTGSGSSAS | 84495.2 | 1.040618935 |
| OD-linker-(GGGGS)3 | 76611.8 | 1.054038875 |
| (GGGGS)3-(GGGGS)5 | 93931.8 | 0.8508051029 |
| GHGTGSTGSGSSAS-(GGGGS)5 | 93161 | 0.857225846 |
| (GGGGS)1-PAPAP | 85626.6 | 0.9180534127 |
| GHGTGSTGSGSSAS-PAPAPAPAPAP | 107703.2 | 0.7178664895 |
| PAPAPAPAPAP-GHGTGSTGSGSSAS | 79214.2 | 0.9751195472 |
| (GGGGS)3-(GGGGS)3 | 79512.2 | 0.9569807011 |
| GSLAEAAAKEAAAKEAAAKAAAAS-PAPAP | 71686.2 | 1.052802305 |
| (GGGGS)5-(GGGGS)5 | 88307.2 | 0.8411495449 |
| PAPAP-GHGTGSTGSGSSAS | 74202 | 0.9973494237 |
| (GGGGS)5-PAPAP | 68637.4 | 1.07169606 |
| (GGGGS)1-GSLAEAAAKEAAAKEAAAKAAAAS | 88444.4 | 0.8135283568 |
| (GGGGS)1-OD-linker | 84654.4 | 0.8341705021 |
| (GGGGS)1-PAPAPAPAPAP | 92285.2 | 0.755394064 |

| | | |
|---|---|---|
| SGLRS-OD-linker | 86897.6 | 0.7903224295 |
| SGLRS-PAPAPAPAPAP | 88440 | 0.7624248199 |
| SGLRS-(GGGGS)5 | 81966.4 | 0.8221808219 |
| OD-linker-PAPAPAPAPAP | 91336.2 | 0.7369481045 |
| (GGGGS)3-GHGTGSTGSGSSAS | 68961.4 | 0.9660291504 |
| PAPAP-GSLAEAAAKEAAAKEAAAKAAAAS | 95794 | 0.684570948 |
| (GGGGS)1-GHGTGSTGSGSSAS | 71963.6 | 0.8986338612 |
| ins119-mScarlet-I3ΔC10 without linkers | 34836.2 | 1.80024562 |
| SGLRS-GHGTGSTGSGSSAS | 69652.4 | 0.8925923931 |
| OD-linker-(GGGGS)5 | 69911.2 | 0.8676497928 |
| GSLAEAAAKEAAAKEAAAKAAAAS-(GGGGS)3 | 60322.8 | 0.9755376132 |
| PAPAPAPAPAP-PAPAPAPAPAP | 88635.2 | 0.6340378595 |
| GSLAEAAAKEAAAKEAAAKAAAAS-GSLAEAAAKEAAAKEAAAKAAAAS | 107739.6 | 0.5089758 |
| PAPAP-(GGGGS)3 | 56712.6 | 0.9548364679 |
| (GGGGS)1-(GGGGS)5 | 62320.4 | 0.8356176796 |
| OD-linker-GSLAEAAAKEAAAKEAAAKAAAAS | 74759.2 | 0.6866624589 |
| PAPAPAPAPAP-(GGGGS)5 | 64734.8 | 0.7864655141 |
| PAPAP-PAPAPAPAPAP | 69174.8 | 0.7180117384 |
| (GGGGS)3-PAPAPAPAPAP | 65651.6 | 0.712257254 |
| PAPAPAPAPAP-(GGGGS)3 | 49421.8 | 0.9421767641 |
| GSLAEAAAKEAAAKEAAAKAAAAS-PAPAPAPAPAP | 69963.4 | 0.6324862803 |
| PAPAPAPAPAP-(GGGGS)1 | 33069 | 1.298884645 |
| PAPAPAPAPAP-GSLAEAAAKEAAAKEAAAKAAAAS | 79638.6 | 0.5344519921 |
| GHGTGSTGSGSSAS-GSLAEAAAKEAAAKEAAAKAAAAS | 63185.2 | 0.6727117811 |
| PAPAPAPAPAP-SGLRS | 33751.2 | 1.238456165 |
| OD-linker-PAPAP | 38943.6 | 1.026287772 |
| (GGGGS)3-(GGGGS)1 | 30510.6 | 1.297617039 |
| GSLAEAAAKEAAAKEAAAKAAAAS-OD-linker | 47216.6 | 0.8306725327 |
| (GGGGS)5-OD-linker | 43920.4 | 0.8535959011 |
| GSLAEAAAKEAAAKEAAAKAAAAS-(GGGGS)5 | 46580.2 | 0.7787838581 |
| mScarlet-I3ΔC10-nnLuz_v5 | 72863.8 | 0.4842615772 |

| | | |
|---|---|---|
| (GGGGS)3-GSLAEAAAKEAAAKEAAAKAAAAS | 51781.2 | 0.6798781066 |
| (GGGGS)5-PAPAPAPAPAP | 50097 | 0.7021284623 |
| OD-linker-(GGGGS)1 | 26949 | 1.202746903 |
| (GGGGS)3-SGLRS | 28411.4 | 1.069992332 |
| GSLAEAAAKEAAAKEAAAKAAAAS-(GGGGS)1 | 23150.4 | 1.270353005 |
| (GGGGS)3-PAPAP | 31090.8 | 0.9303159884 |
| SGLRS-PAPAP | 33428.2 | 0.8385283046 |
| PAPAP-(GGGGS)1 | 22804.4 | 1.164805856 |
| GHGTGSTGSGSSAS-(GGGGS)1 | 20557.6 | 1.21453237 |
| GSLAEAAAKEAAAKEAAAKAAAAS-SGLRS | 19417.2 | 1.193350659 |
| (GGGGS)5-(GGGGS)1 | 18836.6 | 1.190065755 |
| (GGGGS)5-GHGTGSTGSGSSAS | 23020.2 | 0.9588405948 |
| OD-linker-SGLRS | 19049 | 1.155103544 |
| SGLRS-(GGGGS)1 | 21853.8 | 0.9475647139 |
| PAPAP-(GGGGS)5 | 24018.6 | 0.832784541 |
| (GGGGS)1-(GGGGS)1 | 18658.2 | 1.050772485 |
| SGLRS-SGLRS | 22368.6 | 0.8596980469 |
| GHGTGSTGSGSSAS-OD-linker | 22061.4 | 0.8406595962 |
| GHGTGSTGSGSSAS-(GGGGS)3 | 20009.2 | 0.9203736556 |
| PAPAP-SGLRS | 17872 | 1.028478149 |
| PAPAP-OD-linker | 21003.6 | 0.838797182 |
| GHGTGSTGSGSSAS-GHGTGSTGSGSSAS | 19028 | 0.9239291331 |
| (GGGGS)1-SGLRS | 18471.2 | 0.8829483981 |
| GHGTGSTGSGSSAS-SGLRS | 12327.8 | 1.131425069 |
| (GGGGS)1-(GGGGS)3 | 17201.6 | 0.79976212 |
| (GGGGS)5-GSLAEAAAKEAAAKEAAAKAAAAS | 17763.8 | 0.6067947203 |
| SGLRS-(GGGGS)3 | 12176.6 | 0.8006712424 |
| (GGGGS)5-SGLRS | 8659.2 | 1.120528115 |
