## Supplementary Table 4 for "Metabolically robust autoluminescent reporters"

| | Integral luminescence, mean | 593/533 luminescence, mean |
|---|---|---|
| Label on the plot | | |
| nnLuz_v5 | 2768658.8 | 0.4216611596 |
| GHGTGSTGSGSSAS-(GGGGS)1 | 644634 | 1.361064487 |
| OD-linker-(GGGGS)1 | 631970.4 | 1.372132318 |
| (GGGGS)1-GHGTGSTGSGSSAS | 686446.2 | 1.252928776 |
| (GGGGS)1-(GGGGS)1 | 585799.8 | 1.443208981 |
| (GGGGS)1-OD-linker | 704891.2 | 1.192455373 |
| SGLRS-OD-linker | 650647 | 1.277254643 |
| SGLRS-PAPAPAPAPAP | 683420.4 | 1.164427319 |
| (GGGGS)3-SGLRS | 580915.4 | 1.368025156 |
| ins167-tdTomato 2aa linkers | 510393.6444 | 1.534526981 |
| (GGGGS)1-(GGGGS)3 | 607291.6 | 1.285961658 |
| PAPAP-(GGGGS)1 | 556556.8 | 1.365448363 |
| GHGTGSTGSGSSAS-PAPAP | 599166 | 1.267330984 |
| SGLRS-(GGGGS)1 | 482062.4 | 1.534100841 |
| (GGGGS)1-PAPAPAPAPAP | 674203 | 1.092330799 |
| (GGGGS)3-(GGGGS)1 | 531680.6 | 1.371847378 |
| (GGGGS)1-SGLRS | 500292 | 1.440019048 |
| (GGGGS)1-(GGGGS)5 | 602076 | 1.185817367 |
| SGLRS-SGLRS | 472315 | 1.507672032 |
| SGLRS-(GGGGS)3 | 513969.8 | 1.369589269 |
| (GGGGS)5-(GGGGS)1 | 529034.2 | 1.306384234 |
| (GGGGS)1-PAPAP | 517676.4 | 1.332076465 |
| (GGGGS)3-(GGGGS)3 | 547459.4 | 1.206453266 |
| (GGGGS)3-OD-linker | 573526 | 1.144475186 |
| (GGGGS)1-GSLAEAAAKEAAAKEAAAKAAAAS | 651184.25 | 0.999817222 |
| SGLRS-PAPAP | 458807.6 | 1.417086765 |
| OD-linker-(GGGGS)5 | 583063.75 | 1.100738475 |
| SGLRS-GHGTGSTGSGSSAS | 476228 | 1.347195931 |
| (GGGGS)5-(GGGGS)3 | 547570.75 | 1.164448548 |
| GHGTGSTGSGSSAS-SGLRS | 453357.2 | 1.393314815 |

| | | |
|---|---|---|
| GHGTGSTGSGSSAS-(GGGGS)3 | 503389.5 | 1.228510784 |
| (GGGGS)3-GHGTGSTGSGSSAS | 507572.4 | 1.205396392 |
| PAPAP-(GGGGS)3 | 476972.6 | 1.235696248 |
| SGLRS-(GGGGS)5 | 466763.4 | 1.259917333 |
| OD-linker-SGLRS | 419928.8 | 1.389093012 |
| PAPAP-PAPAP | 479108.8 | 1.201769114 |
| GHGTGSTGSGSSAS-PAPAPAPAPAP | 557620.2 | 1.032339018 |
| PAPAPAPAPAP-(GGGGS)1 | 460341.75 | 1.245906355 |
| OD-linker-PAPAPAPAPAP | 562980.2 | 1.016755708 |
| OD-linker-PAPAP | 451681.2 | 1.25357548 |
| PAPAPAPAPAP-SGLRS | 426447 | 1.305326202 |
| GHGTGSTGSGSSAS-(GGGGS)5 | 497469.6 | 1.111582911 |
| OD-linker-(GGGGS)3 | 452340.8 | 1.213436753 |
| (GGGGS)3-PAPAPAPAPAP | 531721.2 | 1.029781968 |
| PAPAP-PAPAPAPAPAP | 527684.25 | 1.030875717 |
| PAPAP-(GGGGS)5 | 484794.2 | 1.121095794 |
| PAPAP-SGLRS | 372227 | 1.444579082 |
| OD-linker-OD-linker | 466815.8 | 1.147039594 |
| (GGGGS)5-(GGGGS)5 | 490562.5 | 1.075050206 |
| GHGTGSTGSGSSAS-GHGTGSTGSGSSAS | 437174.8 | 1.186116922 |
| (GGGGS)3-PAPAP | 429205.8 | 1.19912424 |
| PAPAPAPAPAP-PAPAP | 441752 | 1.158223369 |
| GHGTGSTGSGSSAS-OD-linker | 460871.8 | 1.110005727 |
| SGLRS-GSLAEAAAKEAAAKEAAAKAAAAS | 470438.6 | 1.084102699 |
| OD-linker-GHGTGSTGSGSSAS | 421664 | 1.188547211 |
| (GGGGS)3-(GGGGS)5 | 448973.8 | 1.106171712 |
| PAPAPAPAPAP-PAPAPAPAPAP | 515356.8 | 0.9568556232 |
| PAPAP-GHGTGSTGSGSSAS | 406474.6 | 1.209453015 |
| (GGGGS)5-PAPAP | 407801.4 | 1.189321179 |
| (GGGGS)5-OD-linker | 446433.8 | 1.078647274 |
| (GGGGS)5-SGLRS | 350157 | 1.353330469 |

| | | |
|---|---|---|
| GHGTGSTGSGSSAS-GSLAEAAAKEAAAKEAAAKAAAAS | 509358.8 | 0.9226193117 |
| (GGGGS)5-GHGTGSTGSGSSAS | 406743.8 | 1.131482981 |
| PAPAPAPAPAP-(GGGGS)3 | 401166.2 | 1.137526336 |
| PAPAP-GSLAEAAAKEAAAKEAAAKAAAAS | 457606 | 0.9637762993 |
| PAPAPAPAPAP-(GGGGS)5 | 433245.5 | 1.005328479 |
| (GGGGS)5-PAPAPAPAPAP | 441315.2 | 0.9863868439 |
| PAPAPAPAPAP-GHGTGSTGSGSSAS | 371614 | 1.128619486 |
| (GGGGS)3-GSLAEAAAKEAAAKEAAAKAAAAS | 440365.75 | 0.95216431 |
| GSLAEAAAKEAAAKEAAAKAAAAS-(GGGGS)5 | 415808.6 | 0.9790329812 |
| (GGGGS)5-GSLAEAAAKEAAAKEAAAKAAAAS | 448849.75 | 0.9026101678 |
| PAPAP-OD-linker | 358026.8 | 1.122815375 |
| GSLAEAAAKEAAAKEAAAKAAAAS-(GGGGS)3 | 367291.4 | 1.070299591 |
| GSLAEAAAKEAAAKEAAAKAAAAS-GSLAEAAAKEAAAKEAAAKAAAAS | 463610.2 | 0.8462606055 |
| GSLAEAAAKEAAAKEAAAKAAAAS-PAPAPAPAPAP | 422646.2 | 0.9178205679 |
| GSLAEAAAKEAAAKEAAAKAAAAS-GHGTGSTGSGSSAS | 349412.4 | 1.093006836 |
| GSLAEAAAKEAAAKEAAAKAAAAS-OD-linker | 353640.8 | 1.027607279 |
| PAPAPAPAPAP-OD-linker | 329676.2 | 1.053572719 |
| GSLAEAAAKEAAAKEAAAKAAAAS-PAPAP | 291191.4 | 1.14352413 |
| GSLAEAAAKEAAAKEAAAKAAAAS-(GGGGS)1 | 273329.8 | 1.154420026 |
| PAPAPAPAPAP-GSLAEAAAKEAAAKEAAAKAAAAS | 356322 | 0.8834646975 |
| GSLAEAAAKEAAAKEAAAKAAAAS-SGLRS | 241699 | 1.256296412 |
| OD-linker-GSLAEAAAKEAAAKEAAAKAAAAS | 244241.4 | 0.9299325038 |
