## Supplementary Table 5 for "Metabolically robust autoluminescent reporters"

| | Integral luminescence, me | 608/533 luminescence, mean |
|---|---|---|
| Label on the plot | | |
| nnLuz_v5 ins167-mScarlet-I3ΔC10 without linke | 3615805.9 | 1.80690712 |
| mScarlet3-PAPAP-nnLuz_v5 | 5527632 | 0.784042249 |
| mScarlet3-SGLRS-nnLuz_v5 | 4339185.3 | 0.8691052643 |
| mScarlet3-OD-linker-nnLuz_v5 | 4934250 | 0.7548524321 |
| mScarlet3-(GGGGS)1-nnLuz_v5 | 4354545.1 | 0.8086831794 |
| mScarlet3-GHGTGSTGSGSSAS-nnLuz_v5 | 4561374 | 0.7664722016 |
| mScarlet3-GSLAEAAAKEAAAKEAAAKAAAAS-r | 5220451.5 | 0.6640319315 |
| mScarlet3-(GGGGS)3-nnLuz_v5 | 4512860.7 | 0.7617842161 |
| mScarlet3-(GGGGS)5-nnLuz_v5 | 4611447 | 0.7218296169 |
| mScarlet3-PAPAPAPAPAP-nnLuz_v5 | 4197284.3 | 0.7045053472 |
| nnLuz_v5 ins167-mScarlet-I3 without linkers | 1901025.7 | 1.174018967 |
| nnLuz_v5 ins188-mScarlet-I3 without linkers | 1224109.4 | 1.284470373 |
| nnLuz_v5-GHGTGSTGSGSSAS-mScarlet3 | 1707015.6 | 0.7878991185 |
| nnLuz_v5-(GGGGS)3-mScarlet3 | 1387854.5 | 0.7151333419 |
| nnLuz_v5-(GGGGS)5-mScarlet3 | 1402629 | 0.6950956846 |
| nnLuz_v5-(GGGGS)1-mScarlet3 | 1272092.4 | 0.7027512712 |
| nnLuz_v5-SGLRS-mScarlet3 | 1278276.6 | 0.6897280361 |
| nnLuz_v5-OD-linker-mScarlet3 | 1151785.6 | 0.6736373849 |
| nnLuz_v5-PAPAP-mScarlet3 | 1168734.8 | 0.624195809 |
| nnLuz_v5-PAPAPAPAPAP-mScarlet3 | 1086691.1 | 0.6020983038 |
| mScarlet-I3-SGLRS-nnLuz_v5 | 911927.2 | 0.700013481 |
| nnLuz_v5-GSLAEAAAKEAAAKEAAAKAAAAS-r | 936783.8 | 0.6587056373 |
| nnLuz_v5 ins119 mScarlet3 (2 aa linkers) | 370469.125 | 1.100063837 |
| mScarlet-I3ΔC10-nnLuz_v5 | 504207.4 | 0.6634605355 |
| nnLuz_v5 ins119-mScarlet-I3ΔC10 without linke | 162456.2 | 1.730542979 |
| nnLuz_v5 ins188-mScarlet-I3ΔC10 without linke | 60028.2 | 0.5995087206 |

| | Integral luminescence, mean | 638/533 luminescence, mean |
|---|---|---|
| Label on the plot | | |
| mKate2-GHGTGSTGSGSSAS-nnLuz_v5 | 4832564.9 | 0.3808534447 |
| mKate2-(GGGGS)5-nnLuz_v5 | 5269938.6 | 0.3479379631 |
| mKate2-(GGGGS)1-nnLuz_v5 | 4269504.1 | 0.4176848732 |
| mKate2-GSLAEAAAKEAAAKEAAAKAAAAS-nnLuz_v | 5517365.5 | 0.3183402503 |
| mKate2-PAPAP-nnLuz_v5 | 4267910 | 0.3990619351 |
| mKate2-(GGGGS)3-nnLuz_v5 | 3980646.1 | 0.3694835376 |
| mKate2-OD-linker-nnLuz_v5 | 3921197.9 | 0.3728366751 |
| mKate2-PAPAPAPAPAP-nnLuz_v5 | 4336310.7 | 0.3237362193 |
| nnLuz_v5-GHGTGSTGSGSSAS-mKate2 | 2375677.2 | 0.4989078293 |
| nnLuz_v5 ins188 mKate2 (2 aa linkers) | 830382.275 | 1.265552427 |
| nnLuz_v5-(GGGGS)3-mKate2 | 2164000.2 | 0.4823622524 |
| nnLuz_v5-GSLAEAAAKEAAAKEAAAKAAAAS-mKate | 2236319.2 | 0.4590058165 |
| nnLuz_v5-OD-linker-mKate2 | 2126388.1 | 0.4693117609 |
| nnLuz_v5-PAPAP-mKate2 | 1858476.9 | 0.5145814806 |
| nnLuz_v5-(GGGGS)5-mKate2 | 2166099.5 | 0.4381797394 |
| nnLuz_v5-(GGGGS)1-mKate2 | 2082536.7 | 0.4552922232 |
| mKate2-SGLRS-nnLuz_v5 | 2497987.25 | 0.3693720084 |
| nnLuz_v5-PAPAPAPAPAP-mKate2 | 1892562.3 | 0.4858742305 |
| nnLuz_v5-SGLRS-mKate2 | 1845985.2 | 0.4442225015 |
| nnLuz_v5 ins167-mKate2 without linkers | 1837002.1 | 0.4169868743 |
| nnLuz_v5 ins188-mKate2 without linkers | 755140.2 | 0.9760363945 |
| nnLuz_v5 ins167-mKate2ΔC10 without linkers | 862541.5 | 0.2052889994 |
| mKate2ΔC10-nnLuz_v5 | 439032.6 | 0.1811196811 |
| nnLuz_v5 ins188-mKate2ΔC10 without linkers | 204923 | 0.1751358197 |

| | Integral luminescence, mean | 693/533 luminescence, mean |
|---|---|---|
| Label on the plot | | |
| tdTomato-(GGGGS)1-nnLuz_v5 | 5071149.8 | 1.141169914 |
| tdTomato-SGLRS-nnLuz_v5 | 4437816.7 | 1.213589044 |
| tdTomato-PAPAP-nnLuz_v5 | 4606627.9 | 1.078494763 |
| tdTomato-(GGGGS)3-nnLuz_v5 | 4583048.8 | 0.9925151714 |
| tdTomato-GHGTGSTGSGSSAS-nnLuz_v5 | 4135480.7 | 1.012653189 |
| tdTomato-OD-linker-nnLuz_v5 | 4122851.9 | 1.005627571 |
| tdTomato-PAPAPAPAPAP-nnLuz_v5 | 4134104.1 | 0.9175681187 |
| tdTomato-(GGGGS)5-nnLuz_v5 | 3908536.8 | 0.8680301526 |
| tdTomato-GSLAEAAAKEAAAKEAAAKAAAAS-nnLu | 3389433.9 | 0.8344456005 |
| nnLuz_v5-GHGTGSTGSGSSAS-tdTomato | 1536085.4 | 1.404025727 |
| nnLuz_v5-(GGGGS)5-tdTomato | 1275246.8 | 1.291545498 |
| nnLuz_v5-(GGGGS)3-tdTomato | 1019691.8 | 1.404558741 |
| nnLuz_v5-GSLAEAAAKEAAAKEAAAKAAAAS-tdTo | 1130451.7 | 1.114797249 |
| nnLuz_v5-(GGGGS)1-tdTomato | 887972.3 | 1.362568111 |
| nnLuz_v5-OD-linker-tdTomato | 879805.9 | 1.354938079 |
| nnLuz_v5-SGLRS-tdTomato | 573683.6 | 1.350656165 |
| nnLuz_v5-PAPAP-tdTomato | 590092.4 | 1.184791405 |
